# Modeling Alzheimer’s Disease with APOE4 Neuron-Glial Brain Assembloids Reveals IGFBPs as Therapeutic Targets

**DOI:** 10.1101/2025.10.17.683162

**Authors:** Eliana Sherman, Kevin Qiu, Rebecca Roberts, Lucille Shichman, Sihan Li, Huayu Sun, Lucie Ide, Allison Tucker, Seonjoo Lee, Weronika Gniadzik, Jung-Bum Shin, Katia Sol-Church, Jaideep Kapur, Aiying Zhang, Alev Erisir, Lulu Jiang, the Alzheimer’s Disease Neuroimaging Initiative

## Abstract

Alzheimer’s disease (AD) research has been hindered by the lack of models that faithfully recapitulate the full profile of disease progression in a human genetic background. We developed a 3D assembloid model (“Masteroid”) using iPSC-derived neurons, astrocytes, and microglia from APOE4/4 and isogenic control lines. Neurons were seeded with tau oligomers, then combined with astrocytes and microglia to form mature 3D Masteroids, followed by amyloid-β oligomer exposure. After four weeks, AD-Masteroids exhibited hallmark pathologies, including extracellular amyloid-β deposits, intracellular tau aggregation, neurodegeneration, astrogliosis, and microglial activation, with APOE4 exacerbating all phenotypes. Single-cell RNA sequencing further identified novel roles of IGFBP pathways in amyloid-β and tau-mediated pathology. This innovative platform provides a robust system to dissect cellular and molecular mechanisms of AD progression and offers a powerful tool for therapeutic discovery.

**Highlights:**

- The 3D human neuron–glia assembloid (“Masteroid”), composed of neurons, astrocytes, microglia, and oligodendrocytes, faithfully recapitulates human brain ultrastructure and intercellular interactions.
- Exposure to oligomeric tau and Aβ induced hallmark Alzheimer’s pathologies, including amyloid deposition, tau aggregation, neurodegeneration, and gliosis.
- The APOE4 genotype exacerbated all pathological features, highlighting its role in driving multicellular interactions that accelerate disease progression.
- The IGF signaling axis was identified as a key mediator of Aβ- and tau-induced pathology and a potential therapeutic target.

**Graphical Abstract:** 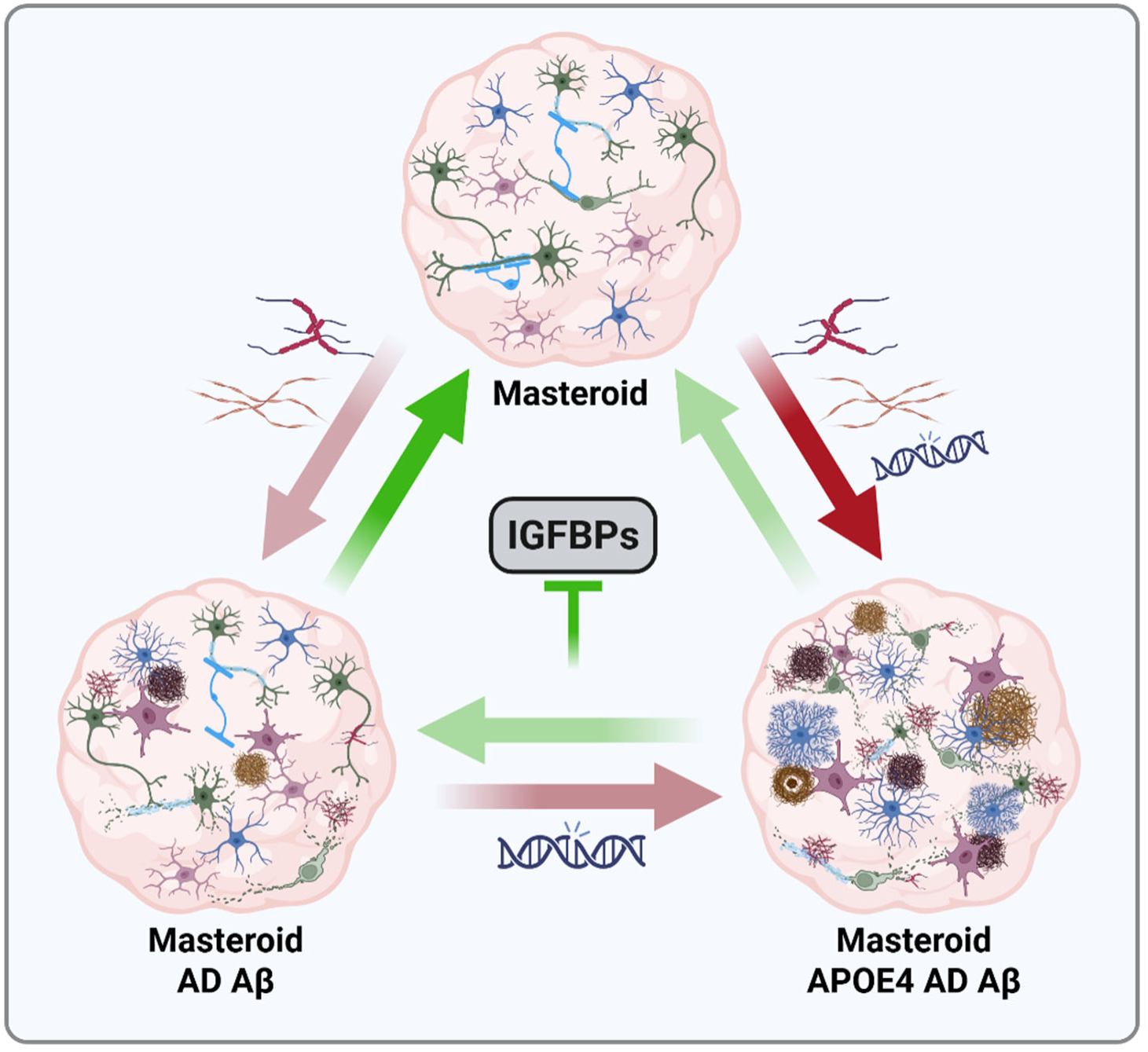

## INTRODUCTION

Alzheimer’s disease (AD), a leading cause of dementia worldwide, is a progressive neurodegenerative disorder that gradually impairs memory, cognition, behavior, and executive functions^1–3^. AD is defined by two primary pathological hallmarks: the accumulation of extracellular amyloid-β (Aβ) plaques and intracellular neurofibrillary tau tangles, accompanied by gliosis and neuronal loss. Despite substantial advances in understanding AD pathology, the complex interplay of genetic, cellular, and environmental factors makes the mechanisms of the disease progression an enigma. Among genetic risk factors, carrying the ε4 allele of apolipoprotein E (APOE4) represents the strongest contributor to late-onset AD^4,5^. Accumulating evidence from epidemiological and genome-wide association studies (GWAS) has shown that APOE4 heterozygotes are two to three times more likely, and homozygotes up to 12 to 15 times more likely, to develop AD compared with APOE3 populations^4,6–8^. In this study, we analyzed positron emission tomography (PET) imaging data from non-demented, preclinical individuals in the Alzheimer’s Disease Neuroimaging Initiative (ADNI) cohort and found that even among cognitively normal control participants, both APOE4 heterozygotes and homozygotes show significantly increased amyloid-β and tau burden compared with non-carriers, demonstrating that APOE4 drives early pathological changes well before clinical symptoms emerge (**Figure 1**). Furthermore, APOE4 increases the risk of amyloid-related imaging abnormalities during treatment with second-generation anti-amyloid monoclonal antibodies and diminishes therapeutic efficacy ^9–11^. Despite its central role in AD pathogenesis, the mechanisms by which APOE4 potentiates pathological amyloid and tau burden, exacerbates disease severity, and confers treatment resistance remain elusive.

**Figure 1.**
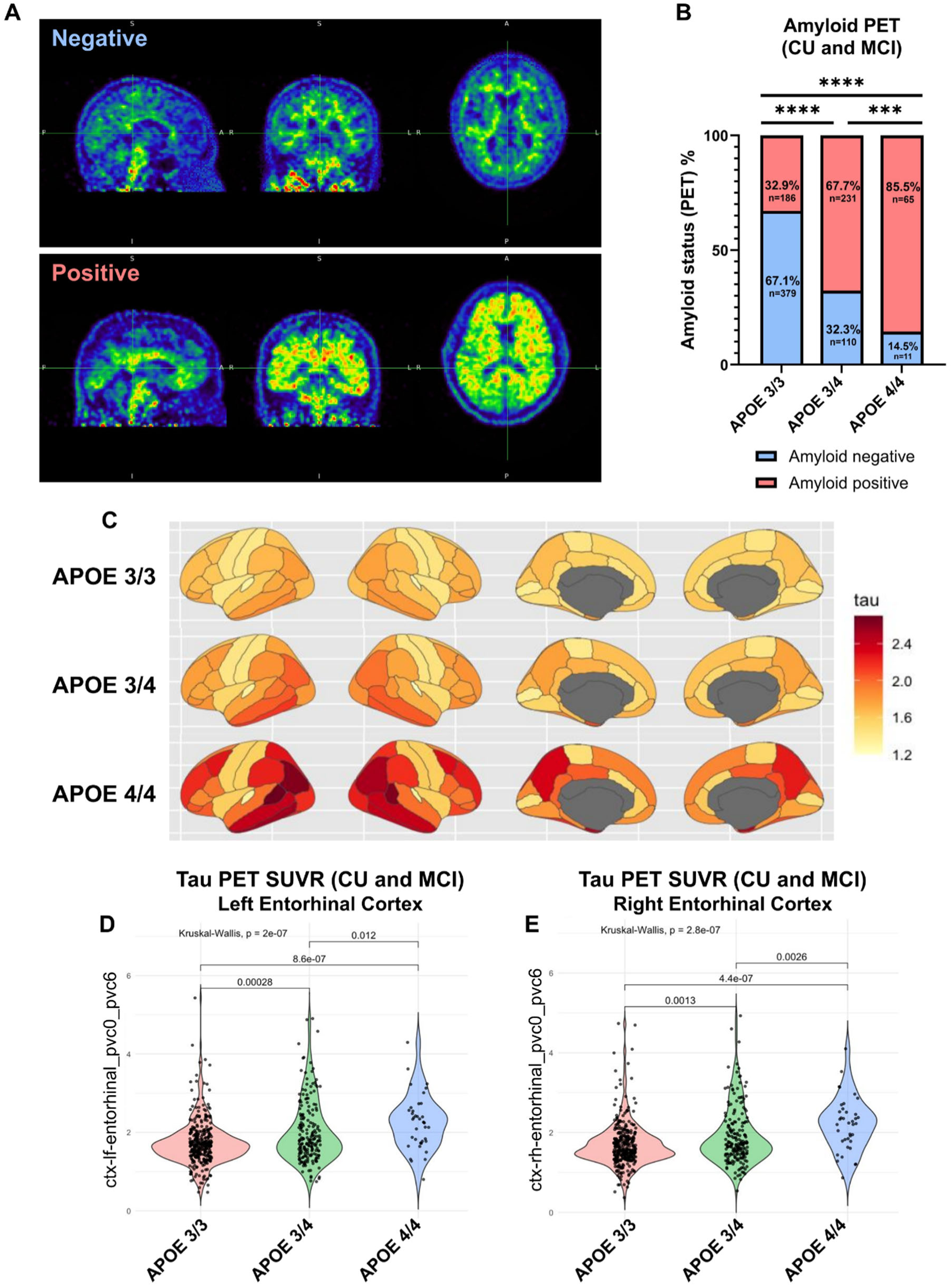
Dose-Dependent Impact of APOE4 on Aβ Deposition and Tau Accumulation from Preclinical to Prodromal Stages. **A.** Examples of a negative (top) and positive (bottom) amyloid PET scan, visualized sagittally (left), coronally (center), and axially (right). A positive scan is defined by a normalized SUVR > 1.11 (see methods). **B.** Quantification of proportion of population with positive amyloid PET scan by APOE genotype in preclinical and prodromal populations (CU and MCI). *n* = 565, 341, and 76 for APOE3/3, APOE3/4, and APOE4/4 respectively. **C.** Mean tau PET burden by brain region in the full cohort, shown by APOE genotypes. **D.** Quantification of normalized Tau PET SUVR in left entorhinal cortex by APOE genotypes in preclinical and prodromal populations (CU and MCI). *n* = 339, 196, and 38 for APOE3/3, APOE3/4, and APOE4/4 respectively. **E.** Quantification of normalized Tau PET SUVR in right entorhinal cortex by APOE genotype in preclinical and prodromal populations (CU and MCI). *n* = 339, 196, and 38 for APOE3/3, APOE3/4, and APOE4/4 respectively. *** = *p* < 0.001, **** = *p* < 0.0001. Amyloid PET statistical analysis was done by Chi-square Test with Yates’ correction (B), and Tau PET statistical analysis was done with Kruskal-Wallis test (D and E) with *p* values shown. See also Figure S1.

Progress in understanding AD and APOE4 has been limited by the lack of models that accurately capture human pathophysiology. While animal models have provided critical insights into amyloid processing, tau aggregation, and neuroinflammation, species-specific differences prevent full recapitulation of the human brain and the complex interplay of AD pathologies. This limitation is particularly pronounced for APOE4, as APOE exhibits species-specific differences in its evolution, structure, and post-translational modifications between rodents and humans^12,13^. Rodent models, even those expressing humanized APOE, encounter difficulty in reproducing the phenotypes observed in human populations^14–16^. Human induced pluripotent stem cell (hiPSC)- based models have emerged as powerful tools to overcome these limitations^17–20^. By reprogramming patient-derived cells into neurons, astrocytes, and other brain cell types, hiPSC models allow the study of AD mechanisms in a human genetic context^21^. Advances in stem cell engineering have further enabled the generation of three-dimensional (3D) brain organoids or assembloids, self-organizing multicellular structures that recapitulate aspects of human brain development, cellular diversity, and architecture^22–26^. Organoids and assembloids provide unique opportunities to investigate cellular interactions and disease processes in a physiologically relevant environment that is difficult to achieve in conventional 2D cultures^27–32^. However, most current organoid systems remain limited in cellular composition, often lacking or exhibiting delayed development of key non-neuronal cell types, and rarely incorporate disease-relevant stressors in a controlled and systematic manner^33–36^.

To overcome the challenges in understanding the pathophysiology of AD and recapitulate human heterogeneity, we built on our previously published Asteroid model^37^, to develop Masteroids, 3D cultures composed of hiPSC-derived neurons, astrocytes, microglia, and endogenously generated oligodendrocytes. In this study, we established Masteroid models using hiPSCs carrying APOE4 and their isogenic APOE3 controls, combining tau and amyloid pathologies to more comprehensively model AD progression. Our Masteroid system was specifically designed to capture both the proteopathic and genetic contributors to AD. Neurons were initially seeded with tau oligomers to prime the system for pathological tau aggregation and then combined with astrocytes and microglia to form mature, multicellular 3D Masteroids. The cultures were subsequently exposed to extracellular amyloid-β oligomers, creating a comprehensive environment that models intermediate and advanced stages of neurodegeneration. After four weeks, AD-Masteroids faithfully recapitulated multiple hallmark features of AD pathology, including Aβ deposition, tau aggregation, widespread neurodegeneration, astrogliosis, and robust microglial activation. Notably, the presence of APOE4 markedly exacerbated all of these pathological phenotypes, highlighting its critical role in disease progression. Electron microscopy revealed that Masteroids recapitulated ultrastructural features of the human brain, including neuronal synaptic structures, neuron-glial interactions, glial phagocytic activity, and early myelination development. Electrophysiological recordings confirmed Masteroid neurons had excitation ability (action potentials in response to current injection) and synaptic events. Leveraging single-cell RNA sequencing, we identified IGFBPs as distinct mediators and highlighted signaling pathways involved in amyloid-β and tau-associated cellular dysfunction, providing new mechanistic insights. Strikingly, treatment of AD-Masteroids with IGFBP inhibitor NBI 31772 significantly reduced neurodegeneration, astrogliosis, and microglial activation, demonstrating a direct role for IGFBP signaling in driving AD pathogenesis. Together, these results establish Masteroids as a powerful tool for high-throughput evaluation of therapeutic interventions, bridging a critical gap between conventional cell cultures and *in vivo* models, and enabling the study of complex genotype-pathology interactions in AD under a controlled environment.

## RESULTS

### Dose-Dependent Impact of APOE4 on Aβ Deposition and Tau Accumulation from Preclinical to Prodromal Stages

AD is composed of three primary stages: preclinical, prodromal, and clinical^38^. In the preclinical stage, patients are primarily asymptomatic and cognitively normal while bearing some pathological markers of AD. In the prodromal stage, patients experience mild cognitive impairment (MCI), ranging from early to late MCI with varying levels of AD pathology. The clinical stage is characterized by extensive AD pathology alongside severe cognitive impairment^38^. Using the ADNI dataset^39–42^, we analyzed the impact of APOE4 status on amyloid and tau burden in human patients during the preclinical and prodromal stages of AD. The ADNI study is composed of three cohorts of varying severities determined by diagnostic criteria of both cognitive and pathological scores: the Cognitively Unimpaired (CU) cohort, the Mild Cognitive Impairment (MCI) cohort, and the Early-Stage Alzheimer’s Disease (DEM) cohort^43,44^. The CU cohort consists of both normal control participants with little to no positive AD pathology as well as cognitively normal participants with some AD pathological markers but little to no cognitive impairment. For our analysis, we considered the CU cohort to be in the preclinical stage, the MCI cohort to be in the prodromal stage, and the DEM cohort to be in the clinical stage.

We began by analyzing the difference in amyloid burden in the preclinical and prodromal stages of AD between APOE3/3, APOE3/4, and APOE4/4 individuals. The amyloid burden was determined by the proportion of a given population with a positive whole brain PET scan which was defined by a normalized Standardized Uptake Value Ratio (SUVR) > 1.11 (**Figure 1A)**. Even one APOE4 allele markedly increased amyloid positivity in preclinical and prodromal patients, with homozygotes showing the highest rates, indicating a dose-dependent effect of APOE4 on disease progression (**Figure 1B**). APOE3/4 in the CU group showed significantly higher amyloid positivity than non-carriers, though no difference was observed between APOE3/4 and APOE4/4 carriers (**Figure S1A**). By MCI stage, the APOE4/4 individuals had a significantly greater proportion of amyloid positive scans than the APOE3/4, though this difference is not significant in the DEM cohort as nearly all APOE3/4 and APOE4/4 patients had positive amyloid scans. In both the MCI and DEM cohorts, any instance of APOE4 (APOE4/4 vs APOE4/3 vs APOE3/3) led to a significantly greater proportion of amyloid positive scans than in non-carriers (**Figures S1B and S1C**). Across all genotypes, each additional APOE4 allele was associated with higher amyloid positivity, with APOE4/4 carriers showing the greatest amyloid burden (**Figure S1D**).

Amyloid accumulation typically marks early AD pathology while tau burden increases in parallel with cognitive decline^45^. We therefore analyzed tau PET data in the same ADNI cohort by APOE status across disease stages, focusing on the entorhinal cortex, which is among the first regions to show tau pathology^46,47^. Tau burden was determined by the actual SUVR value instead of a binary cutoff of positive or negative scan like amyloid prevalence (**Figure 1C**). Similarly to observations with amyloid burden, APOE4 heterozygotes and homozygotes had significantly greater tau burden than non-carriers in both the left and right entorhinal cortex as early as in the preclinical and prodromal stages of AD with APOE4 homozygotes exhibiting greater tau burden than APOE4 heterozygotes (**Figures 1D and 1E**). In the CU cohort, tau burden in the left entorhinal cortex did not differ significantly by APOE4 status, though a strong trend was seen between APOE3/3 and APOE4/4. In the right entorhinal cortex, tau burden was significantly higher in APOE4/4 than APOE3/3 individuals, with no difference between APOE3/3 and APOE3/4 (**Figure S1E**). By the prodromal stage of MCI, APOE4 heterozygotes and homozygotes showed significantly greater tau burden in both the left and right entorhinal cortices compared to non-carriers (**Figure S1F**). Across all genotypes, each additional APOE4 allele was linked to higher tau burden in both entorhinal cortices, with APOE4 homozygotes showing the greatest levels (**Figure S1G**).

Altogether, our analysis demonstrated a dose-dependent effect of the APOE4 allele on disease progression, influencing both Aβ deposition and tau accumulation from the earliest, preclinical, and prodromal stages.

### 3D Masteroid culture system recapitulates human brain cell types and ultrastructure

To establish a 3D neuron-glial assembloid model that recapitulates the coordinated origin and emergence of neurons, astrocytes, and microglia, similar to human brain development, we developed a 3D coculture system, termed Masteroid, for modeling neurodegenerative diseases, following a modified neuron-astrocyte AstTau assembloid protocol developed in our previous work ^37,48^. In brief, human iPSCs with a normal APOE3/3 genotype were initially differentiated into hematopoietic progenitor cells and neural progenitor cells (hiNPCs) where hiNPCs were also further differentiated into astrocyte progenitor cells. All three types of progenitor cells were subsequently differentiated into neuronal, astrocytic, and microglial lineages to generate human iPSC-derived neurons (hiNeurons), astrocytes (hiAstrocytes), and microglia (hiMicroglia). These cells were then combined into a 3D spherical culture, termed Masteroids, and allowed to mature over the course of 5 weeks (**Figure 2A**). We also maintained individual hiNeurons, hiAstrocytes, and hiMicroglia in 2D culture for 1 week alongside the creation of Masteroids to confirm proper cell morphology and identity through immunolabeling (**Figure 2B**). Mature Masteroids exhibited extensive neuronal and astrocytic processes, with microglia distributed throughout the Masteroid (**Figure 2C**).

**Figure 2.**
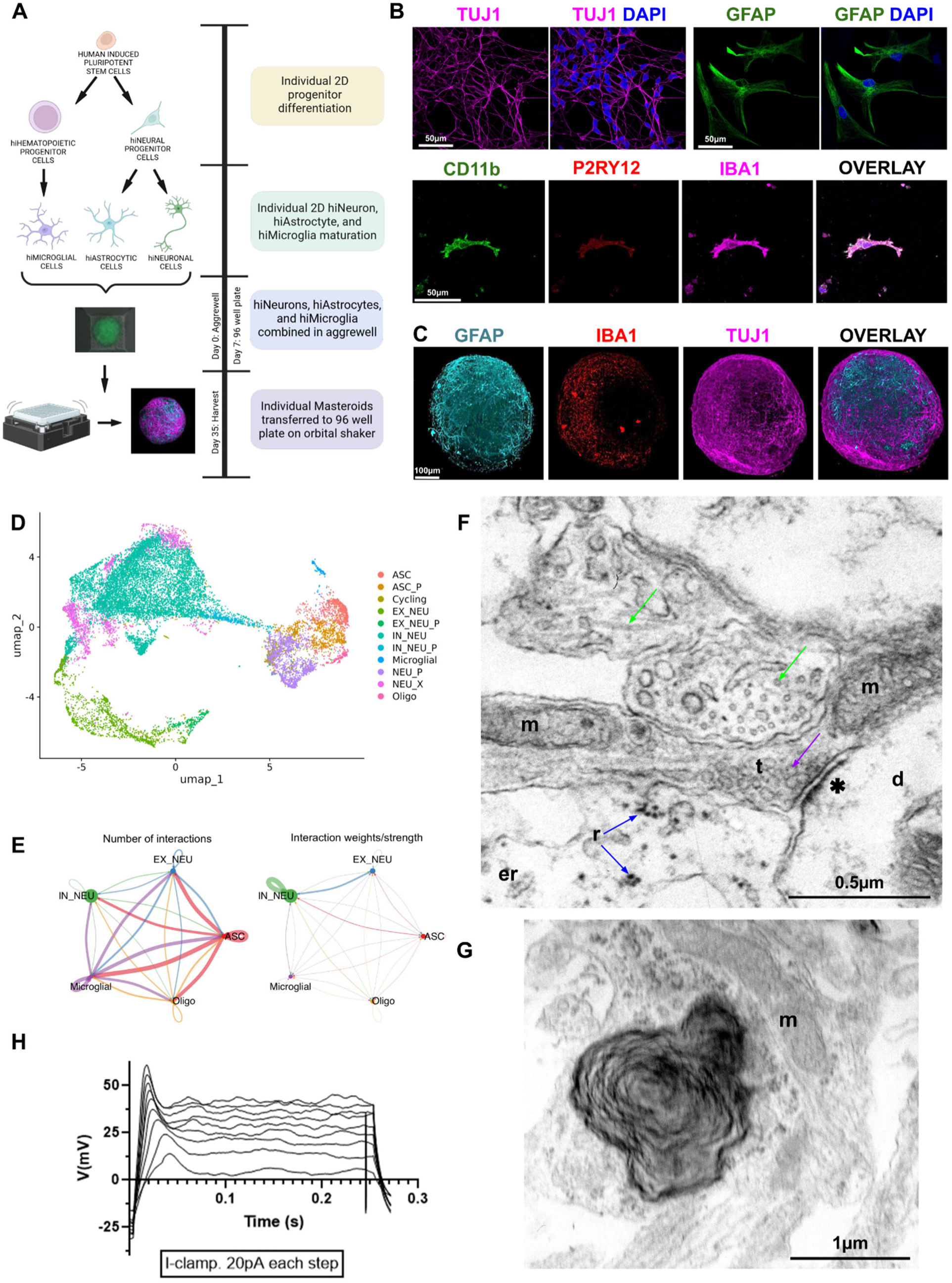
3D Masteroid culture system recapitulates human brain cell types and ultrastructure. **A.** Schematic illustrating experimental Masteroid culture system. Created with BioRender.com. **B.** Representative immunofluorescence images of iPSC-derived neurons (TUJ1, magenta), astrocytes (GFAP, green), and microglia (CD11b, green; P2RY12, red; and IBA1, magenta) with dsDNA stain Dapi (blue). Scale bar = 50µm. **C.** Representative tile scanned and z-stacked immunofluorescence images of 3D Masteroid with cell type markers for astrocytes (GFAP, cyan), microglia (IBA1, red), and neurons (TUJ1, magenta). Scale bar = 100µm. **D.** UMAP of single-cell transcriptomic profiles by scRNA-seq after filtration and cell type identification. The following cell types were identified: astrocytes (ASC, red), excitatory neurons (EX_NEU, lime green), inhibitory neurons (IN_NEU, teal), microglia (Microglial, blue), oligodendrocytes (Oligo, pink), and various progenitor and pan-neuronal cells (ASC_P, orange; Cycling, olive; EX_NEU_P, green; IN_NEU_P, cyan; NEU_P, purple; NEU_X, light pink). **E.** Number and weight of significant ligand-receptor interactions between five major cell types. Circle sizes are proportional to the number of cells within a given cluster, and line width is proportional to the indicated number or strength of interactions. Cell types included are excitatory neurons (EX_NEU, blue), inhibitory neurons (IN_NEU, green), astrocytes (ASC, red), microglia (Microglial, purple), and oligodendrocytes (Oligo, orange). **F.** Electron microscopy (EM) image of a synapse within a Masteroid. Green arrows point to microtubules, purple arrow points to synaptic vesicles, blue arrows point to ribosomes, and ✱ indicates postsynaptic density. Scale bar = 0.5µm. **G.** EM image of a myelin whirl within a Masteroid. Scale bar = 1µm. **H.** Single cell patch-clamp (I-clamp) recording of an action potential in a Masteroid in response to current injection (20 pA per step). (F-G) m: mitochondria, t: synaptic terminal, d: dendrite, er: endoplasmic reticulum, and r: ribosomes. See also Figure S2.

The final Masteroid cellular phenotypes were characterized using single-cell RNA sequencing (scRNA-seq), profiling 8,000 cells per sample at a sequencing depth of 50,000 read pairs per cell. Visualization by uniform manifold approximation and projection (UMAP) demonstrated the separation of cells into trajectory guided clusters. The clusters were identified as astrocytes (ASC), excitatory neurons (EX_NEU), inhibitory neurons (IN_NEU), microglia (Microglial), oligodendrocytes (Oligo), and various progenitor and pan-neuronal cells (**Figure 2D**). Analysis of scRNA-seq clustering revealed an oligodendrocyte cluster, which was confirmed by immunolabeling for Myelin Basic Protein (MBP)–positive myelinating oligodendrocytes in Masteroids (**Figures 2D, S2A, and S2B**). To determine the origin of the oligodendrocytes, the astrocyte, oligodendrocyte, and progenitor clusters were isolated and redefined on a UMAP followed by a pseudotime subclustering and trajectory analysis (**Figure S2C**)^49–51^. This analysis traced gene expression trajectories and indicated that oligodendrocytes in Masteroids likely originate from astrocyte progenitor cells. Previous studies showed that astrocytes and oligodendrocytes share a common glial precursor^52–54^, and astrocytes can be reprogrammed into oligodendrocytes^55,56^. Together with our pseudo-time analysis, this supports that the endogenously generated oligodendrocytes within our Masteroids originate from astrocyte progenitor cells. To determine the number and probability of intercellular interactions between cell types, we did CellChat scRNA-seq analysis which identified the differentially over-expressed ligands and receptors in each cluster^57^. Significant receptor-ligand interactions were present between all cell types in the Masteroids, noting the presence of extensive intercellular communication, with the strongest interaction probability occurring between the neuronal clusters (**Figure 2E**). scRNA-seq profiling of voltage-gated channel expression in hiNeurons confirmed Masteroid functionality, showing expression of hyperpolarization cyclic-nucleotide gated (HCN) subunits for I_h_ currents, voltage-gated sodium and potassium channels for action potentials, and calcium channel subunits required for synaptic neurotransmitter release (**Figures S2D, S2E, S2F, and S2G**)^58–60^.

To further validate that Masteroids recapitulate the ultrastructure of human brain tissue, we performed transmission electron microscopy (EM). Synapses were identified (**Figure 2F**), with presynaptic terminals containing numerous vesicles adjacent to the synaptic cleft and a prominent postsynaptic density, features indicative of active synapses and supporting the biophysical functionality of the Masteroid model ^61,62^. EM also revealed the proper ultrastructural morphology of mature neuronal and glial cells within Masteroids (**Figures S2H, S2I, and S2J**). Masteroid neurons have abundant perinuclear cytoplasm with mitochondria, endoplasmic reticulum, Golgi, and numerous ribosomes, and large, round nuclei with low-contrast, unclustered chromatin and distinct nucleoli (**Figure S2H**)^63,64^. Masteroid neurons often extend large neurites from the cell body (**Figure S2H**)^64^. Masteroid glial cells display irregularly-shaped and darker nuclei, composed of dense, clumped chromatin with smaller and darker cytoplasm (**Figure S2I**). Cells with intracellular space containing a full ensemble of cytoskeletal elements including microtubules and neurofilaments were also found (**Figure S2J**)^61,65^. Myelin whirls, concentric membrane stacks lacking a neuropil core, were observed, confirming myelinating oligodendrocyte development in Masteroids (**Figure 2G**)^63^. Moreover, whole-cell patch-clamp recordings further validated the electrophysiological functionality of whole Masteroids at day 35 of culture (**Figure S2K**). Masteroid hiNeurons exhibited a resting membrane potential, and action potentials were evoked by current injection (**Figures S2L and 2H**)^66^. The hiNeurons also showed synaptic transmission with relatively infrequent but consistent and spontaneous excitatory postsynaptic currents (EPSCs) observed, indicating the formation of synapses between hiNeurons within the Masteroids, as was also observed by EM **(Figures S2M and 2F**).

Together, scRNA-seq profiling, EM ultrastructural analysis, and electrophysiological recordings demonstrate the overall biophysical functionality, cellular health, and intercellular connectivity of Masteroids.

### Propagation of oTau and Aβ oligomers in Masteroids mirrors key features of AD

We next used our Masteroid system to build a model of sporadic AD by selectively propagating human oligomeric tau (oTau) and/or Aβ oligomers (**Figure S3A**). Oligomeric tau was extracted from post-mortem human brain tissue by centrifugation using a well-established extraction method^67–70^. Human oTau fractions were isolated from diseased Alzheimer’s brains (AD oTau) and age matched control brains (NC) and identified by Western blot with anti-Tau antibodies (total by Tau-5 and Phospho-Tau by pThr217) (**Figures S3B and S3C).** The dose-response of oTau toxicity was determined using LDH analysis at 24 and 96 hours after addition of oTau in 2D hiNeurons (**Figure S3D**). Based on the dose-response, 0.04mg/mL of human oTau was seeded to hiNeurons 24 hours prior to their combination with hiAstrocytes and hiMicroglia for all AD-Masteroids while the controls were treated with the same volume of NC fraction. Purified Aβ oligomers (0.5µM/mL) were spiked in at one week after the Masteroids were transferred to 96 well plates. The proper size (<10 kDa) and identity of the purified Aβ oligomers was confirmed by western blot with anti-Aβ antibody (6E10) (**Figure S3E**). To dissect the contributions of amyloid and tau pathology to AD pathogenesis, we established four Masteroid conditions, NC, NC Aβ, AD oTau, and AD Aβ, by selectively introducing NC or AD oTau together with either Aβ oligomers or a DMSO vehicle (**Figure 3A**). Subsequent analysis of these Masteroids revealed significant development of AD-associated pathology in all diseased conditions, with the most pronounced effects observed in Masteroids treated with both AD oTau and Aβ oligomers (**Figure 3**).

**Figure 3.**
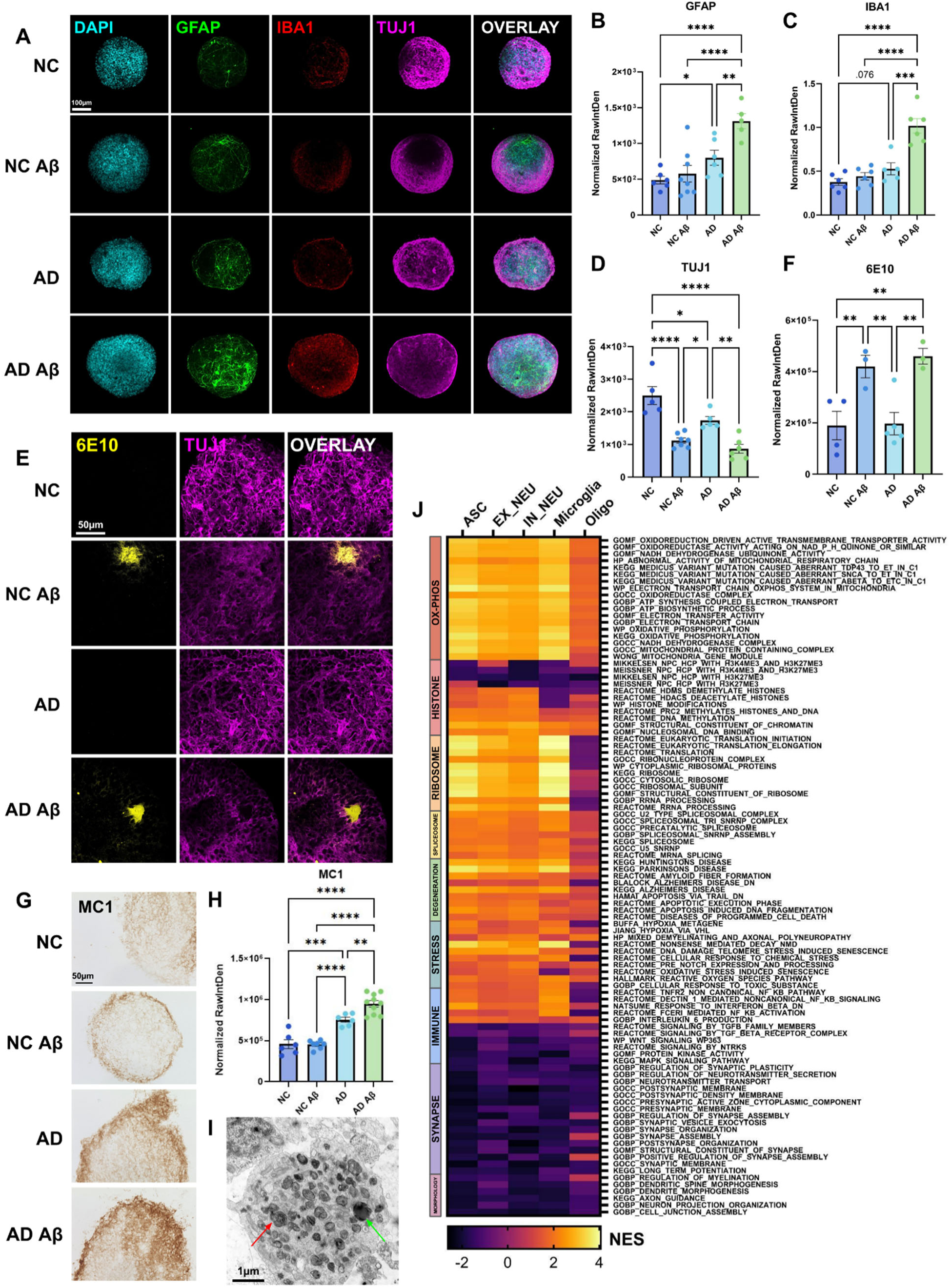
Propagation of oTau and Aβ oligomers in Masteroids mirrors key features of AD. **A.** Tiled and z-stacked immunofluorescence images of control (NC), Aꞵ oligomers only (NC Aꞵ), oTau only (AD), and Aꞵ oligomers and oTau (AD Aꞵ) Masteroids with dsDNA nuclei stain (DAPI, cyan), astrocytes (GFAP, green), microglia (IBA1, red), and neurons (TUJ1, magenta). Scale bar = 100µm. **B.** Quantification of astrogliosis by GFAP raw integrated density normalized to Masteroid area. **C.** Quantification of microglia activation by IBA1 raw integrated density normalized to TUJ1. **D.** Quantification of neuronal loss by TUJ1 raw integrated density normalized to Masteroid area. **E.** Immunofluorescence images of Aꞵ plaques (6E10, yellow). Scale bar = 50µm. **F.** Quantification of Aꞵ burden by 6E10 raw integrated density normalized to Masteroid area. **G.** DAB staining images of misfolded tau (MC1). **H.** Quantification of misfolded tau by MC1 DAB staining normalized to Masteroid area. **I.** Electron microscopic view of neurodegenerative pathology in AD oTau Masteroid: dystrophic neurite containing degenerated mitochondria (red arrow), multilamellar whorls, and electron dense granules (green arrow). Scale bar = 1µm. **J.** Heatmap showing normalized enrichment scores (NES) of pathways in APOE3 AD Aβ Masteroids versus APOE3 NC Masteroids by scRNA-seq. Positive NES indicates upregulation in APOE3 AD Aβ. Pathways include oxidative phosphorylation (dark orange), histone modifications (pink), ribosomal (light orange), spliceosomal (yellow), neurodegenerative (green), stress response (teal), immune signaling (blue), synaptic (purple), and neuronal/glial morphology (burgundy). Error bars indicate 95% of the standard error of the mean. * = *p* < 0.05, ** = *p* < 0.01, *** = *p* < 0.001, **** = *p* < 0.0001. Statistical analysis was done using one-way ANOVA with Fisher’s LSD (B, C, D, F) and Tukey’s multiple comparison test (H). See also Figures S3 and S4.

The reactive astrogliosis marker GFAP showed significant increases in astrocytic reactivity in both AD oTau and AD Aβ Masteroids, with a more significant increase in the AD Aβ condition compared to AD oTau alone (**Figure 3B**). While AD oTau Masteroids exhibited some microglial activation as indicated by IBA1, only AD Aβ Masteroids showed significant increase (**Figure 3C**). Both Aβ and AD oTau treatments led to a significant decrease in neuronal TUJ1 immunolabeling compared to NC Masteroids, with Aβ conditions showing greater neuronal loss than AD oTau alone, and the most severe loss observed in AD Aβ Masteroids (**Figure 3D**). Immunolabeling by 6E10 in whole Masteroids revealed that both NC Aβ and AD Aβ conditions formed Aβ plaques (**Figures 3E and 3F**). While some plaques were more diffuse, others exhibited the classic “star” shape morphology with a dense core^71^. No Aβ plaques were observed in the NC or AD oTau conditions. In identifying the tau pathology, we found that the AD oTau and AD Aβ Masteroids showed significant tau misfolding (MC1 antibody) with the AD Aβ Masteroids containing significantly more misfolded tau than the AD oTau only condition (**Figures 3G and 3H**). This is consistent with the most pronounced glial activation and neuronal loss occurring in the AD Aβ condition, highlighting the deleterious synergistic interaction between tauopathy and Aβ in AD^72,73^. Interestingly, there were no significant differences in neuronal loss or Aβ burden between NC Aβ and AD Aβ Masteroids (**Figures 3D, 3E and 3F**), suggesting that tau may not directly induce or enhance amyloid pathology. EM imaging also revealed extensive neurodegenerative and AD pathology in NC Aβ, AD oTau, and AD Aβ Masteroids, including abundant dystrophic neurites, degenerating myelin, membrane sheath disintegration, glial engulfment, and lipofuscin accumulation (**Figures 3I, S3F, S3G, S3H, S3I, and S3J**)^74–76^.

To further delineate cell type-specific responses to amyloid and tau pathology, we compared scRNA-seq profiles from NC, AD oTau, NC Aβ, and AD Aβ Masteroids, uncovering both shared and distinct transcriptional alterations induced by oTau and Aβ. UMAP projections demonstrated successful integration of all scRNA-seq datasets and visualized changes in the relative proportions of cell types across conditions (**Figures S4A, S4B, S4C, and S4D**). Differential gene expression analysis of the IN_NEU and EX_NEU clusters revealed an overall reduction in synaptic gene expression in AD Aβ Masteroids compared to NC Masteroids (**Figures S3K and S3L**), and ASC clusters exhibited disease-associated astrocyte features, such as extensive upregulation of PLCG2 in AD Aβ Masteroids (**Figure S3M**)^77,78^. Additionally, the predicted interaction probability between EX_NEU and IN_NEU clusters was notably decreased in NC Aβ and AD Aβ Masteroids relative to NC, consistent with observed neuronal loss by immunolabeling and the reduction in synaptic markers (**Figures S4I, S4J, S4K, S4L, S4Q, S4R, S4S, and S4T**). Functional gene set enrichment analysis (GSEA) further revealed that AD Aβ Masteroids robustly recapitulate AD-related pathway alterations (**Figure 3J**). They showed strong enrichment in respirasomal, spliceosomal, ribosomal, neurodegenerative, and stress-response pathways, alongside marked dysregulation of histone and immune signaling^79^. Conversely, pathways linked to neuronal and glial morphology, synaptic formation, and function were substantially downregulated, consistent with neuronal loss and impaired communication^79,80^. Overall, these results highlight the strong modeling of AD-associated cellular dysfunction in AD Aβ Masteroids.

### oTau and Aβ Oligomer Seeding in APOE4/4 Masteroids Reveals APOE4-Genotypic AD Pathology

APOE4 Masteroids were generated using hiNeurons, hiAstrocytes, and hiMicroglia derived from homozygous APOE4 iPSCs and treated with the same selective seeding of oTau and Aβ oligomers as APOE3/3 Masteroids, as described in **Figure 3**. Cell morphology and identity were confirmed through individual 2D cultures and immunostaining of whole Masteroids (**Figures 4A and 4B**). Proper neuronal and glial ultrastructure within APOE4 Masteroids was validated by EM imaging (**Figures S5A and S5B**), which revealed synaptic vesicles, extensive cytoskeletal organization with microtubules, neurofilaments, and dendritic branching (**Figures S5C, S5D, S5E**). Extensive processes were also observed by both IF and EM imaging (**Figures 4B and S5F**). Cellular phenotypes of APOE4 Masteroids were further characterized by scRNA-seq, with UMAP projections demonstrating integration across experimental conditions and visualizing relative proportional shifts in cell types between groups (**Figures S4E, S4F, S4G, and S4H**).

**Figure 4.**
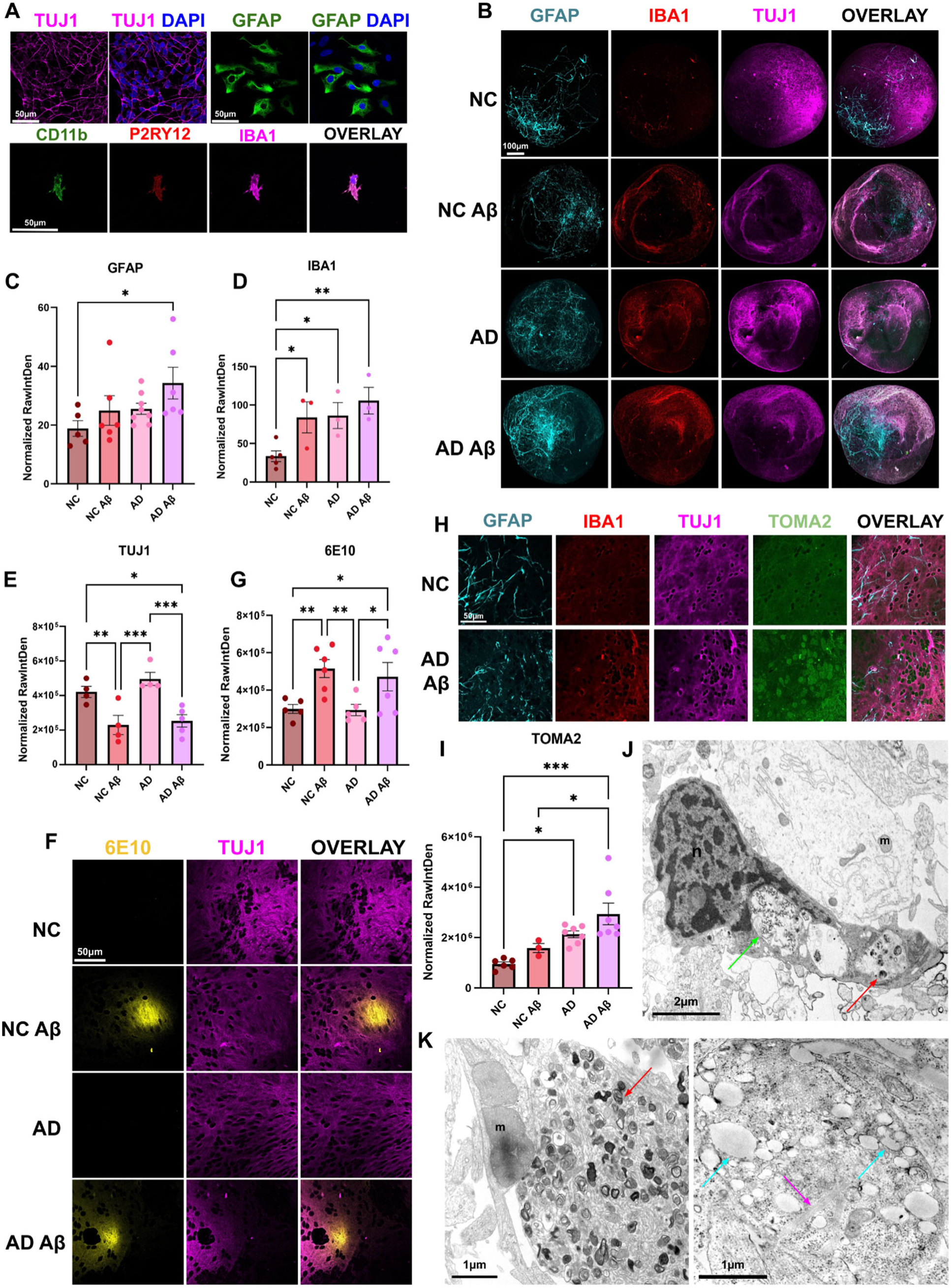
oTau and Aβ Oligomer Seeding in APOE4/4 Masteroids Reveals APOE4-Genotypic AD Pathology. **A.** Immunofluorescence images of APOE4 iPSC induced neurons (TUJ1, magenta), astrocytes (GFAP, cyan), and Microglia (CD11b, green; P2RY12, red; and IBA1, magenta) with dsDNA stain DAPI. Scale bar = 50µm **B.** Tile scanned and z-stacked immunofluorescence images of control (NC), Aꞵ oligomer only (NC Aꞵ), oTau only (AD), and Aꞵ oligomer and oTau (AD Aꞵ) APOE4 Masteroids with astrocytes (GFAP, green), microglia (IBA1, red), and neurons (TUJ1, magenta). Scale bar = 100µm. **C.** Quantification of astrogliosis by GFAP labeling, raw integrated density normalized to Masteroid area. **D.** Quantification of microglial activation by IBA1 labeling, raw integrated density normalized to Masteroid area. **E.** Quantification of neuronal loss by TUJ1 labeling, raw integrated density normalized to Masteroid area. **F.** Immunofluorescence images of amyloid plaques (6E10, yellow) within Masteroids. Scale bar = 50µm. **G.** Quantification of amyloid burden by 6E10 labeling, raw integrated density normalized to Masteroid area. **H.** Immunofluorescence images of NC and AD Aꞵ APOE4 Masteroids with astrocytes (GFAP, cyan), microglia (IBA1, red), neurons (TUJ1, magenta), and oligomeric tau (TOMA2, green). Scale bar = 50µm. **I.** Quantitation of oligomeric tau labeled by TOMA2 antibody, raw integrated density normalized to neuronal area. **J.** Electron microscopy (EM) image of glial engulfment in an APOE4 AD Aꞵ Masteroid. Green arrow points to loose chromatin engulfed while the red arrow points to engulfed dystrophic neurites. Scale bar = 2µm. **K.** EM images of neurodegeneration in APOE4 AD (left) and APOE4 AD Aꞵ (right) Masteroids. Red arrow points to dystrophic neurites and degenerating myelin, “m” depicts deformed and enlarged mitochondria, cyan arrows depict lysosomes, and magenta arrow points to filament bundles. Scale bars = 1µm. (J-K) n: nucleus, m: mitochondria. Error bars indicate 95% standard error of the mean. * = *p* < 0.05, ** = *p* < 0.01, *** = *p* < 0.001, **** = *p* < 0.0001. Statistical analysis was done using one-way ANOVA with Fisher’s LSD (C, D, E, G) and Tukey’s multiple comparisons test (I). See also Figures S4 and S5.

Pathological analyses were performed on APOE4 Masteroids to determine the impact of oTau and Aβ in this deleterious genotype. The reactive astrocyte marker GFAP showed a significant increase in astrogliosis between APOE4 NC and APOE4 AD Aβ Masteroids, confirmed by both immunofluorescence labeling and immunoblotting (**Figures 4C, S5G, and S5I**). scRNA-seq analysis revealed elevated reactive astrocyte gene expression in the APOE4 AD Aβ ASC population compared to APOE4 NC ASC, with significant upregulation of AD risk genes PLCG2 and CLU and enrichment of post-mortem AD astrocyte signatures (**Figure S5M and S5N**) ^81–85^. Significant microglial activation was observed across all non-control APOE4 Masteroid conditions, with the strongest activation in APOE4 AD Aβ Masteroids (**Figures 4D, S5H, and S5J**). scRNA-seq analysis demonstrated robust enrichment of disease-associated microglial (DAM) signatures, with key genes including NEAT1, PICALM, APOE, MAPT, CD44, PSEN1, SORL1, and GRN markedly upregulated in APOE4 AD Aβ Masteroids, closely mirroring post-mortem AD brain profiles (**Figures S5O and S5P**)^84–87^. Notably, PLCG2 showed striking upregulation, and IGF1R emerged as one of the most prominently elevated transcripts in APOE4 AD Aβ Masteroids (**Figure S5O**). Neuronal analyses revealed significant loss in APOE4 NC Aβ and AD Aβ Masteroids compared to both APOE4 NC and AD oTau conditions (**Figures 4E, S5H, and S5K**). scRNA-seq profiling further showed an increase in both inhibitory and excitatory neurons carrying AD-associated transcriptional signatures in APOE4 AD Aβ Masteroids compared to APOE4 NC (**Figures S5Q, S5R, S5S, and S5T**)^82^. Interestingly, no differences were detected between APOE4 NC and AD oTau, or between APOE4 NC Aβ and AD Aβ groups in neuronal loss as indicated by TUJ1 intensity (**Figure 4E**).

Whole-organoid immunolabeling of APOE4 Masteroids with the 6E10 antibody revealed extensive Aβ plaque accumulation in both NC Aβ and AD Aβ conditions (**Figures 4F and 4G**). Notably, despite the absence of exogenous Aβ oligomers, 6E10-positive plaques were detected in non-Aβ APOE4 Masteroids. Although plaques were observed only in the APOE4 NC condition and not in the APOE4 AD oTau condition by whole-organoid labeling, both NC and AD oTau conditions exhibited plaque formation in 14μm cryosections (**Figures S5U and S5V**), indicating that the APOE4 genotype alone is sufficient to drive endogenous plaque formation. APOE4 Masteroids also developed extensive tau pathology, with significant increases in hyperphosphorylated (labeled by pThr217, pSer202), oligomeric (labeled by TOMA2), and misfolded tau (MC1) in AD oTau and AD Aβ conditions (**Figures 4H, 4I, S5G, S5L, S5W, S5X, S5Y, and S5Z**). The presence of Aβ did not further affect tau hyperphosphorylation on site pThr217, as NC and NC Aβ, and AD oTau and AD Aβ groups, showed comparable levels within each category (**Figures S5G, S5L, S5W, and S5X**). However, tau oligomerization and misfolding, assessed with TOMA2 and MC1 antibodies, followed a more severe pattern: AD oTau and AD Aβ Masteroids developed significantly greater pathology than NC and NC Aβ groups, and the addition of Aβ oligomers exacerbated these effects. APOE4 AD Aβ Masteroids developed the highest burden of oligomeric and misfolded tau, surpassing AD oTau Masteroids (**Figures 4I, S5Y, and S5Z**). Moreover, ultrastructural analyses by EM revealed extensive neurodegenerative pathology. Dystrophic neurites, lipofuscin, and degenerating myelin were detected across all conditions, including APOE4 NC Masteroids (**Figures 4K, S5AA, S5BB, and S5CC**). Morphologically abnormal mitochondria and pronounced vacuole accumulation were particularly evident in AD and AD Aβ Masteroids (**Figures 4K, S5CC, and S5DD**). In addition, APOE4 AD Aβ Masteroids displayed striking glial engulfment, with cytoplasmic inclusions containing phagocytosed material, ranging from loose chromatin to dystrophic neurite fragments (**Figures 4J and S5EE**).

Altogether, APOE4 Masteroids alone developed Aβ plaques, mild gliosis, and neuronal dystrophy, whereas addition of oTau and Aβ oligomers induced robust AD-like pathology, including reactive astrocyte and microglial activation, neuronal loss, Aβ plaques, tau hyperphosphorylation, oligomerization and misfolding, and ultrastructural neurodegeneration, with the greatest severity observed in APOE4 AD Aβ Masteroids.

### APOE4/4 significantly exacerbates AD pathology compared to APOE3/3

Having determined the impact of AD pathology within genotypes, we wanted to examine the impact of the APOE4 genotype on AD pathology in comparison to the APOE3 isogenic control. Notably, APOE4 NC Masteroids develop Aβ plaques even in the absence of Aβ oligomer or AD oTau challenge, which differs from APOE3 Masteroids (**Figures S5U and S5V**). In APOE4 Masteroids, neuronal loss did not differ significantly between NC and AD oTau conditions, whereas APOE3 AD oTau Masteroids exhibited a significant reduction relative to APOE3 NC (**Figures 3D and 4E**). However, when directly comparing between the genotypes, APOE4 Masteroids exhibited significantly more neuronal loss compared to their APOE3 isogenic control with the most profound difference observed in the basal NC conditions (**Figures 5A and 5B**). This suggested that the lack of significant difference observed across the intermediate APOE4 conditions was due to an increase in severity at the basal level of APOE4 rather than a lack of neuronal loss in APOE4 AD oTau Masteroids. This is further evidenced by the significant increase in neuronal loss in APOE4 AD oTau and AD Aβ Masteroids when directly compared to their respective APOE3 counterparts (**Figure 5B**). Additionally, APOE4 Aβ Masteroids had significantly greater Aβ burdens than APOE3 Aβ Masteroids, and the plaques in APOE4 Aβ Masteroids were generally larger and denser than those in APOE3 Aβ Masteroids (**Figures 5A and 5C**). There was also a remarkable increase in misfolded tau in APOE4 Masteroids compared to APOE3 Masteroids, particularly in the NC and NC Aβ conditions (**Figures 5D and 5E**).

**Figure 5.**
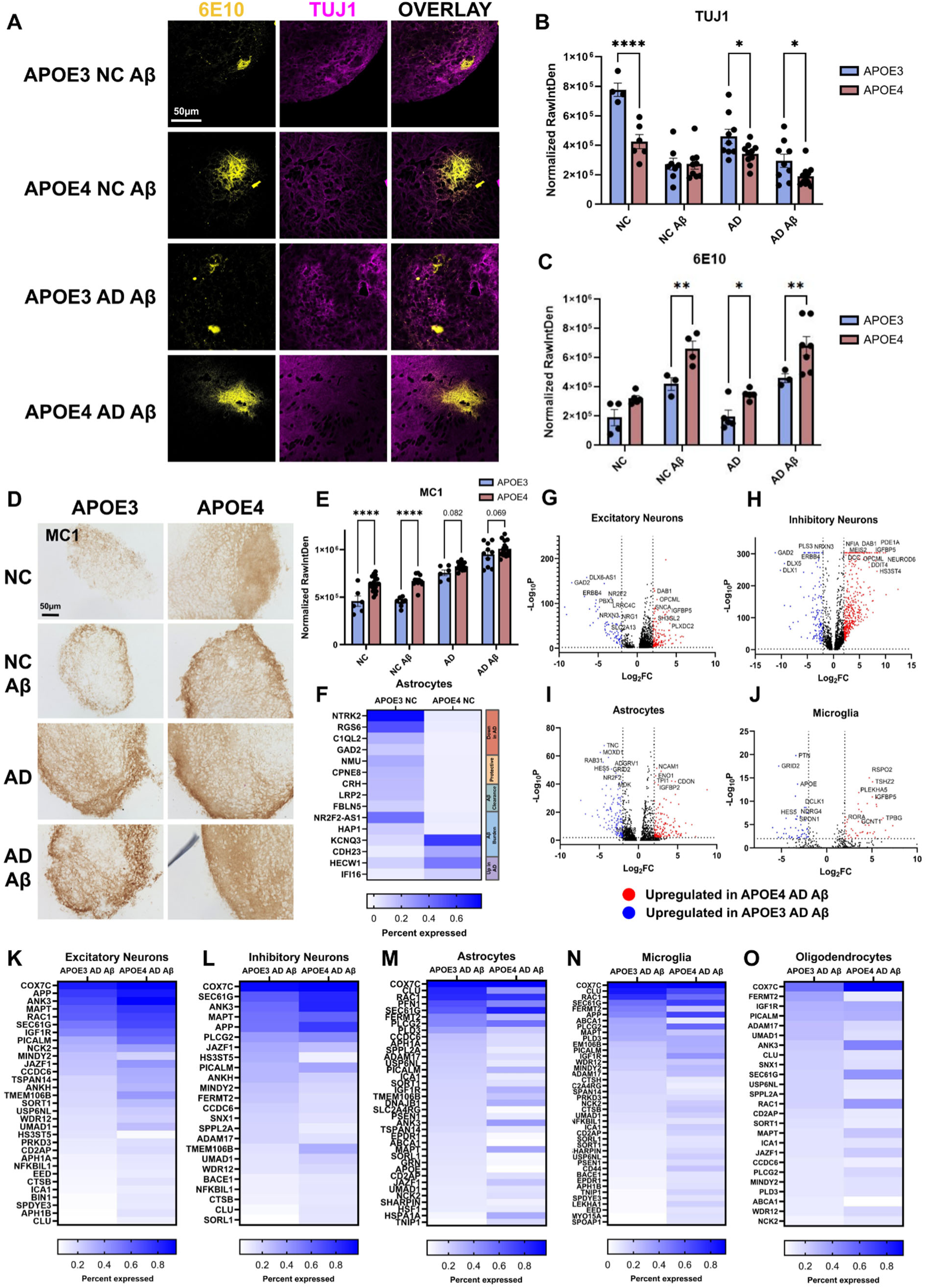
APOE4/4 significantly exacerbates AD pathology compared to APOE3/3. **A.** Immunofluorescence images of amyloid plaques (6E10, yellow) within Aβ Masteroid conditions. Scale bar = 50µm. **B.** Quantification of neuronal loss by TUJ1 labeling, raw integrated density normalized to Masteroid area. **C.** Quantification of Aβ plaques by 6E10 labeling, raw integrated density normalized to Masteroid area. **D.** Representative images of MC1 in 14µm thin sections of APOE3 and APOE4 Masteroids. Scale bar = 50µm. **E.** Quantification of misfolded tau in APOE3 and APOE4 Masteroids by MC1 raw integrated density normalized to Masteroid area. **F.** Expression of top DEGs (LFC > 3) of AD-associated genes in APOE3 NC and APOE4 NC astrocytes (ASC) by scRNA-seq. Genes are color-coded as follows: downregulated in AD (red), protective against AD (yellow), involved in Aβ clearance (green), contributing to Aβ burden (blue), or upregulated in AD (purple). **G-J:** Red dots indicate genes upregulated in APOE4, and those in blue are downregulated in APOE4. Log fold-change cutoffs are y = −2 and 2, with a *p* value cutoff of 0.05 (-log10P > 2). **G.** Volcano plot of DEGs between APOE3 AD Aꞵ and APOE4 AD Aꞵ excitatory neurons (EX_NEU). **H.** Volcano plot of DEGs between APOE3 AD Aꞵ and APOE4 AD Aꞵ inhibitory neurons (IN_NEU). **I.** Volcano plot of DEGs between APOE3 AD Aꞵ and APOE4 AD Aꞵ astrocytes (ASC). **J.** Volcano plot of DEGs between APOE3 AD Aꞵ and APOE4 AD Aꞵ microglia (Microglial). **K.** Expression of differentially expressed AD-associated genes in APOE3 AD Aβ and APOE4 AD Aβ excitatory neurons by scRNA (EX_NEU). **L.** Expression of differentially expressed AD-associated genes in APOE3 AD Aβ and APOE4 AD Aβ inhibitory neurons by scRNA (IN_NEU). **M.** Expression of differentially expressed AD-associated genes in APOE3 AD Aβ and APOE4 AD Aβ astrocytes by scRNA (ASC). **N.** Expression of differentially expressed AD-associated genes in APOE3 AD Aβ and APOE4 AD Aβ microglia by scRNA (Microglial). **O.** Expression of differentially expressed AD-associated genes in APOE3 AD Aβ and APOE4 AD Aβ oligodendrocytes by scRNA (Oligo). Error bars indicate 95% standard error of the mean. * = *p* < 0.05, ** = *p* < 0.01, *** = *p* < 0.001, **** = *p* < 0.0001. Statistical analysis was done using two-way ANOVA with Fisher’s LSD (B, C, and E). Blue bars are APOE3 conditions, burgundy bars are APOE4 conditions. See also Figures S4 and S6.

Comparison of glial pathology between APOE4 and APOE3 Masteroids revealed that APOE4 Masteroids exhibit a more severe pathological baseline. In APOE3 Masteroids, significant astrogliosis was observed under intermediate conditions and reached its peak in the AD Aβ condition. By contrast, in APOE4 Masteroids, only the AD Aβ condition showed significantly greater astrogliosis relative to the APOE4 baseline (**Figures 3B and 4C**). This was further supported by scRNA-seq profiling of several top DEGs (LFC > 3) between APOE3 NC and APOE4 NC astrocytic clusters (**Figure 5F**). APOE4 NC astrocytes showed reduced expression of protective and amyloid-clearance genes (NTRK2, C1QL2, GAD2, NMU, LRP2, HAP1) and upregulation of genes linked to Aβ accumulation and neuroinflammation (CDH23, IFI16)^88–94^, indicating a transcriptomic shift toward a more pathogenic state (**Figure 5F**). Notably, many of the top 20 up- and downregulated DEGs across major cell types in APOE3 NC versus APOE4 NC Masteroids were linked to AD, indicating widespread dysregulation of AD-associated genes driven by the APOE4 genotype alone (**Figures S6A, S6B, S6C, S6D, and S6E**)^84^. In APOE3 Masteroids, only the most severe condition combining AD oTau and Aβ oligomers induced significant microglial inflammation. However, all non-control APOE4 conditions exhibited significant microglial activation, with the strongest response observed in the AD Aβ condition (**Figures 3C and 4D**). This trend suggests that microglial activation required less pathology in APOE4

Masteroids than in APOE3 Masteroids, indicating that APOE4 itself primes the Masteroids for an amplified inflammatory response. scRNA-seq profiling further highlighted genotype- and cell type–specific dysregulation of AD-associated genes. PLCG2 expression was elevated at baseline in APOE4 NC and remained higher across all cell types under AD Aβ challenge, consistent with a primed inflammatory state in APOE4 Masteroids (**Figure S6F**). CLU expression also diverged by genotype: APOE3 astrocytes and microglia upregulated CLU under AD Aβ, whereas APOE4 microglia downregulated CLU and APOE4 neurons aberrantly increased it, indicating a dysregulated Clusterin response in APOE4 Masteroids (**Figure S6G**)^95,96^.

To further determine the impact of APOE4 in pathologically challenged conditions, we narrowed the scRNA-seq analysis to compare between APOE4 AD Aβ and APOE3 AD Aβ, which revealed widespread genotype- and cell type-specific dysregulation of AD-associated genes (**Figures 5G, 5H, 5I, 5J, and S6H**). In excitatory (EX_NEU) and inhibitory (IN_NEU) neurons, APOE4 Masteroids showed strong upregulation of APP, MAPT, PICALM, TMEM106B, CLU, BACE1, and SORT1, while astrocytes (ASC) and microglia (MG) exhibited selective increases in MAPT, CLU, APOE, PSEN1, and PLCG2, and oligodendrocytes (Oligo) showed similar trends for PICALM, MAPT, PLCG2, and SORT1 (**Figures 5K, 5L, 5M, 5N, and 5O**). CellChat network analysis revealed enhanced neuronal-glial interaction probabilities in APOE4 compared to APOE3 (**Figure S6I**), indicating a more reactive and pathologically engaged cellular network. By cross-referencing a scRNA-seq atlas of the entorhinal cortex from AD brains, our cluster-level analyses revealed stronger AD-associated neuronal and microglial signatures in APOE4 AD Aβ Masteroids, accompanied by downregulation of protective A2 astrocyte signatures (**Figures S6J, S6K, S6L, and S6M**)^82,97^. Collectively, these results demonstrate that APOE4 amplifies AD-relevant transcriptional dysregulation and intercellular network alterations under Aβ and oTau-induced stress, producing a more severe, genotype-dependent pathology. APOE4 drives stronger, networked AD pathology than APOE3, underscoring its central role in genotype-dependent disease progression.

### APOE4, Aβ oligomers, and oTau synergistically activate AD-related pathways, highlighting IGF signaling as a key mediator

To delineate how APOE4 interacts with Aβ and oTau to influence AD pathogenesis, we performed Cytoscape STRING network analysis of differentially expressed genes (DEGs) across all major cell types in APOE4 AD Aβ Masteroids relative to APOE3 NC. Network analysis revealed extensive transcriptional reprogramming across cell types. Excitatory neurons (EX_NEU) and inhibitory neurons (IN_NEU) exhibited dysregulation of translational and ribosomal stress, metabolic pathways, synaptic activity, chaperone-mediated protein folding, and cytoskeletal signaling (RHO GTPase in EX_NEU; ubiquitination and proteolysis in IN_NEU) (**Figures 6A and 6B**). Inhibitory neurons (IN_NEU) showed further dysregulation in neurogenesis and cholesterol biosynthesis (**Figures 6A**). Astrocytes (ASC) showed enrichment for translational and ribosomal stress, mitochondrial dysfunction, neurodegeneration-related pathways, metabolism, neurogenesis and development, cell-matrix interactions and synaptic activity, and immune/inflammatory responses (**Figures 6C**). Microglia (Microglial) exhibited transcriptional changes in translation, immune and inflammatory responses, proteolysis, migration, axon guidance, and Wnt signaling (**Figures 6D**)^98,99^. Oligodendrocytes (Oligo) were enriched for ribosome and translation, transcriptional signaling, cell adhesion and synapse formation, oxidative phosphorylation and neurodegeneration pathways, development, and DNA damage responses (**Figures 6E**).

**Figure 6.**
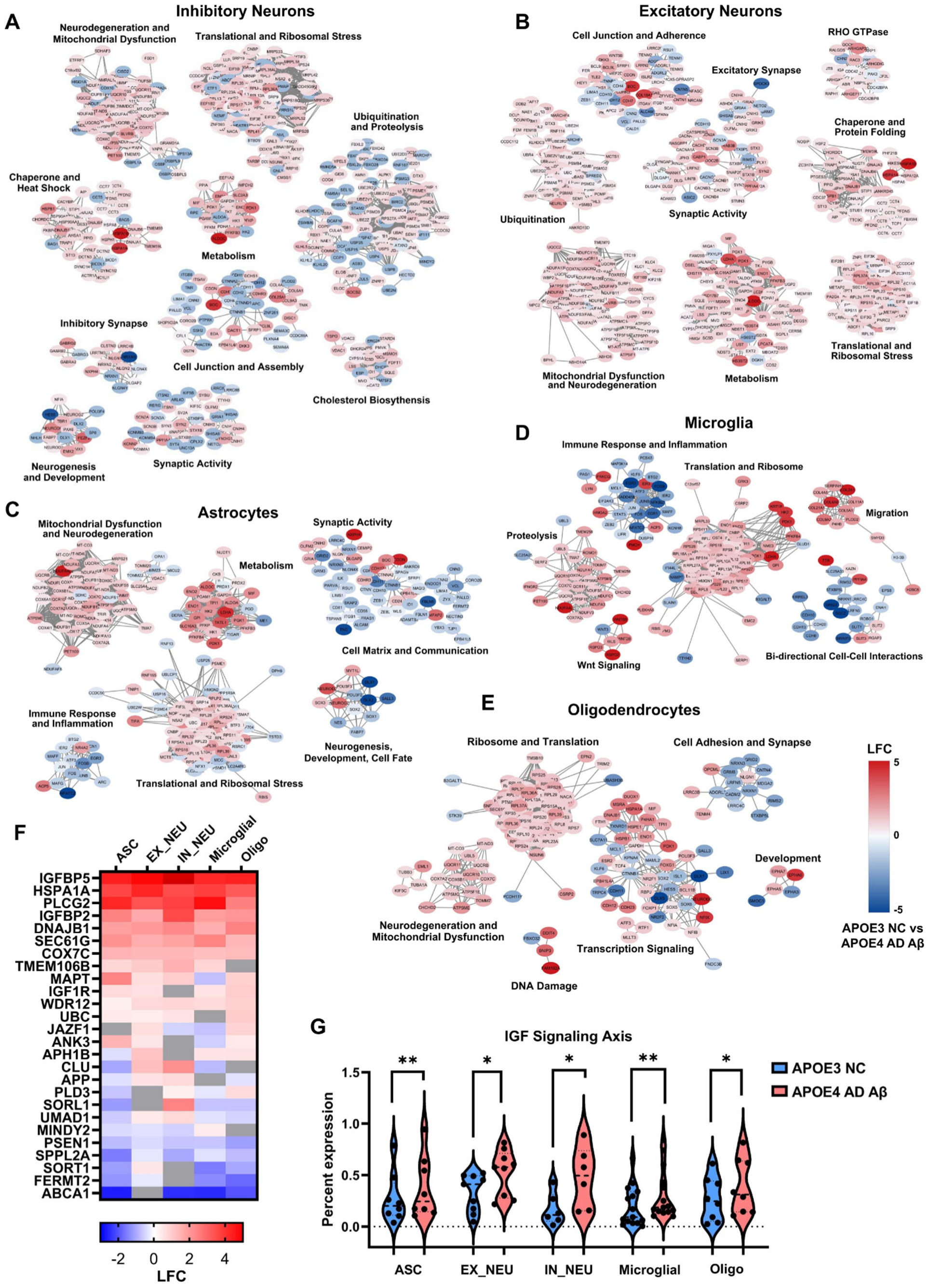
APOE4, Aβ oligomers, and oTau synergistically activate AD-related pathways, highlighting IGF signaling as a key mediator. **A-E:** MCL clustering of DEGs between APOE3 NC and APOE4 AD Aβ cell types (see Methods, *p* < 0.05) with additive gene annotations. Genes upregulated in APOE4 AD Aβ are shown in red, and those downregulated are shown in blue. **A.** Gene interaction network of significant DEGs in inhibitory neurons (IN_NEU). **B.** Gene interaction network of significant DEGs in excitatory neurons (EX_NEU). **C.** Gene interaction network of significant DEGs in astrocytes (ASC). **D.** Gene interaction network of significant DEGs in microglia (Microglial). **E.** Gene interaction network of significant DEGs in oligodendrocytes (Oligo). **F.** scRNA-seq heatmap of log fold changes of AD-associated genes between APOE3 NC and APOE4 AD Aβ. Genes upregulated in APOE4 AD Aβ are shown in red, and those downregulated are shown in blue. **G.** scRNA-seq comparison of IGF signaling gene expression in astrocytes (ASC), excitatory neurons (EX_NEU), inhibitory neurons (IN_NEU), microglia (Microglial), and oligodendrocytes (Oligo) between APOE3 NC and APOE4 AD Aβ Masteroids. * = *p* < 0.05, ** = *p* < 0.01. Statistical analysis was done using paired one-sided Wilcoxon signed-rank test (G). See also Figure S6.

Next, analysis of top differentially expressed transcripts in APOE4 AD Aβ versus APOE3 NC highlights APOE4’s broad impact on multicellular AD-relevant networks, with IGFBP2, IGFBP5 and IGF1R, key components of the IGF signaling pathway, among the most significantly altered genes (**Figures 6F and S6N**). Across all cell types, DEGs implicated in the IGF signaling axis were prominently altered, suggesting APOE4 modulates neurotrophic and metabolic networks in response to Aβ and oTau stress (**Figures 6G, S6O, S6P, S6Q, S6R, and S6S**)^100^. Collectively, these results demonstrate that APOE4 synergizes with Aβ and oTau to drive widespread, cell type– specific transcriptional networks that converge on protein homeostasis, synaptic function, metabolism, and neuroinflammation, reinforcing its central role in exacerbating AD pathology.

### IGFBP inhibitor NBI 31772 ameliorates AD-associated neuroinflammation

Our scRNA-seq data suggested that Insulin-like Growth Factor Binding Proteins (IGFBPs) and the Insulin-like Growth Factor (IGF) signaling pathway play a crucial role in the propagation of AD pathology in the Masteroid model (**Figures 5G, 5H, 5I, 5J, S6H, 6F, and 6G**). The IGF axis is a complex signaling pathway increasingly implicated in AD pathogenesis^100–108^. IGFBPs, particularly IGFBP2 and IGFBP5, have been shown to modulate IGF availability, neuronal survival, and synaptic function^101,102,109^. To test the involvement of IGFBPs and the IGF axis in AD pathological development, we treated Masteroids with an IGFBP inhibitor, a small molecule named NBI 31772, that has been successfully applied in retinal organoids previously^109^. All eight Masteroid conditions (±APOE4, ± oTau, ± Aβ oligomers) were treated with NBI 31772 at 0.5µM for 4 days to assess the role of IGF signaling in AD pathology. Treatment markedly reduced reactive astrogliosis, neuronal loss, and microglial inflammation in diseased Masteroids (**Figures 7 and S7**). In APOE3 NC Aβ Masteroids, NBI 31772 decreased astrogliosis but did not restore neuronal loss, whereas APOE3 AD oTau Masteroids showed no change in astrogliosis but exhibited increased TUJ1 labeling, indicating reduced neuronal loss (**Figures 7A, 7B, and 7C**). In the severe APOE3 AD Aβ condition, NBI 31772 ameliorated both astrogliosis and neuronal loss (**Figures 7B and 7C**). Similar effects were observed in APOE4 Masteroids: treated NC Aβ, AD oTau, and AD Aβ Masteroids all showed significantly less neuronal loss than untreated controls, though improvements in astrogliosis and microglial inflammation were limited to the severe APOE4 AD Aβ condition, with intermediate conditions largely unaffected (**Figures 7B, 7D, 7E, 7F, S7A, S7B, S7C, S7D, S7E, S7G, and S7H**). While more APOE4 conditions showed partial neuronal recovery upon NBI 31772 treatment, the effect was more pronounced in APOE3 Masteroids (**Figure 7B**). This likely reflects the higher baseline neuronal loss in APOE4 NC, APOE4 AD oTau, and APOE4 AD Aβ Masteroids, where immunolabeling was already substantially reduced compared to APOE3 counterparts (**Figure 5B**). Nevertheless, in all conditions exhibiting neuronal recovery, NBI 31772 restored neurons to the baseline levels typical of their respective genotypes, indicating that treatment effectively normalized neuronal capacity relative to the intrinsic genotype-specific baseline (**Figure 7B**).

**Figure 7:**
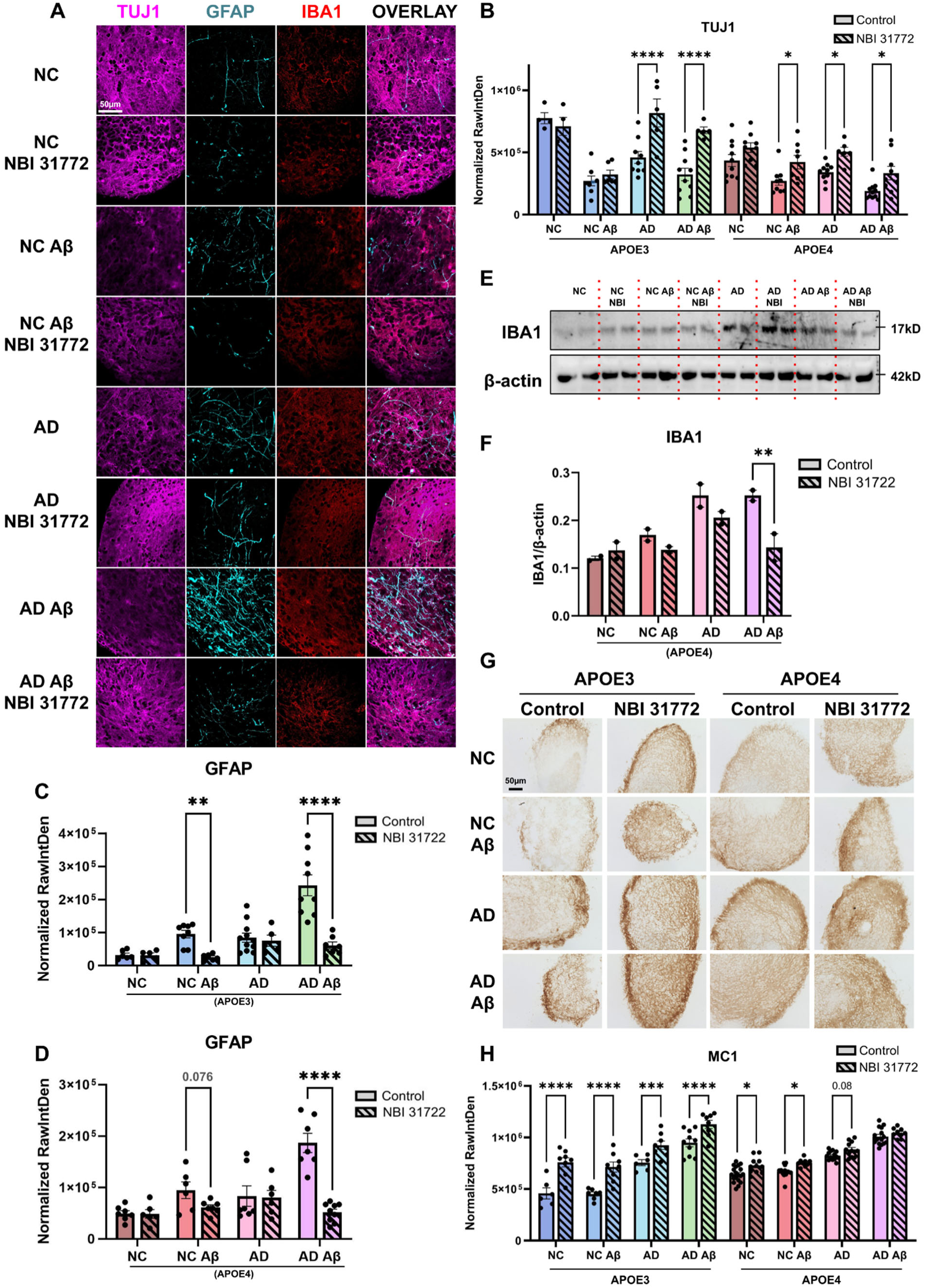
IGFBP inhibitor NBI 31772 ameliorates AD-associated neuroinflammation. **A.** Immunofluorescence images of APOE3 Masteroids ± NBI 31772 treatment. Neurons (TUJ1, magenta), astrocytes (GFAP, cyan), and microglia (IBA1, red) are shown. Scale bar = 50µm. **B.** Quantification of neuronal loss in all conditions by TUJ1 raw integrated density normalized to Masteroid area. **C.** Quantification of astrogliosis in APOE3 Masteroids by GFAP raw integrated density normalized by Masteroid area. **D.** Quantification of astrogliosis in APOE4 Masteroids by GFAP raw integrated density normalized by Masteroid area. **E.** Western blot band of IBA1 and corresponding β-actin. **F.** Quantification of microglial activation in APOE4 Masteroids by IBA1 Western blot raw integrated density with background subtraction normalized by β-actin. **G.** DAB images of MC1 in 14µm thin APOE3 and APOE4 Masteroid sections ± NBI 31772 treatment. Scale bar = 50µm. **H.** Quantification of misfolded tau burden in Masteroids ± NBI 31772 treatment by MC1 raw integrated density normalized to Masteroid area. Error bars indicate 95% standard error of the mean. * = *p* < 0.05, ** = *p* < 0.01, *** = *p* < 0.001, **** = *p* < 0.0001. Statistical analysis was done using two-way ANOVA with Fisher’s LSD (B, C, D, E, and G). Bars with stripes represent NBI 31772 treatment. See also Figure S7.

Surprisingly, despite the pronounced reduction in glial activation and neuronal loss, NBI 31772 treatment led to an increase in tau phosphorylation and misfolding (**Figures 7G, 7H, S7F, S7H, S7I, and S7J**). Tau misfolding (MC1) increased markedly in all APOE3 conditions following treatment, whereas only APOE4 NC and NC Aβ Masteroids showed significant increases. Consistent with the less pronounced neuronal recovery, the increase in tau misfolding was less substantial in APOE4 Masteroids compared to APOE3 Masteroids (**Figures 7G and 7H**). Moreover, NBI 31772 treated APOE4 NC and NC Aβ Masteroids exhibited higher tau hyperphosphorylation at pThr217 in 14µm sections, while APOE4 NC and AD Aβ Masteroids showed increased pSer202 phosphorylation by immunoblot (**Figures S7F, S7H, S7I, and S7J**).

Overall, NBI 31772–mediated inhibition of IGFBPs mitigates AD-associated neuroinflammation and neuronal loss across APOE3 and APOE4 Masteroids, while paradoxically increasing tau phosphorylation and misfolding, highlighting a complex, genotype-dependent role of IGF signaling in AD pathology.

## DISCUSSION

Alzheimer’s disease (AD) is a complex neurodegenerative disorder in which the interplay between genetic risk factors and pathological hallmarks such as amyloid-beta (Aβ) and tau drives disease progression^1,110^. Modeling these interactions in a human-relevant system has remained challenging, particularly in the preclinical and prodromal stages of AD. Our APOE3 and APOE4 Masteroid model provides a robust platform to interrogate both individual components and their synergistic effects on AD pathology, while also enabling functional studies of genotype-specific contributions and therapeutic interventions.

Consistent with clinical observations, our analysis with the pan ADNI PET scan imaging dataset revealed that APOE4 markedly exacerbates amyloid deposition and tau pathology in preclinical and prodromal stages, with severity increasing with allele dosage^111^. Notably, in our Masteroid model, APOE4 NC organoids developed Aβ plaques and tau hyperphosphorylation even in the absence of exogenous seeding of Aβ oligomers or oTau, highlighting the intrinsic pathogenic influence of the APOE4 genotype. These findings recapitulate the early, genotype-driven disease processes that are nearly impossible to study in living humans. By incorporating excitatory and inhibitory neurons, astrocytes, microglia, and oligodendrocytes, our Masteroid brain organoid system matures rapidly within five weeks while maintaining functional synapses, proper membrane potential, action potential propagation, and active ligand-receptor interactions. The endogenously generated oligodendrocytes in the Masteroids further provides the potential to study myelin dynamics, neurodegeneration, and other CNS disorders, including multiple sclerosis, in a versatile and reproducible human-relevant system^112^. The functional robustness of our Masteroids is extensively underscored by electrophysiological recordings, which demonstrate healthy, well-formed neuronal networks capable of patch-clamp analysis. This confirms both the biophysical integrity and the intercellular connectivity necessary for modeling complex AD pathology. Compared to traditional cerebral organoids or rodent models^15,28,29,33^, our system achieves functional maturity and multi-lineage composition in a fraction of the time, bridging a critical gap in translational research.

At the molecular level, scRNA-seq reveals that APOE4 significantly amplifies AD-associated transcriptional programs across neurons and glia. By combining oTau and Aβ in our assembloids, a feature largely absent in previous APOE4 organoid or spheroid models, we provide the first system capable of dissecting the combined impact of multiple pathological components in a human genotype-specific context. Importantly, our data demonstrates that APOE4 not only exacerbates neurodegeneration and glial activation but also modulates critical pathways including the IGF signaling axis^105^, highlighting mechanisms that may underlie genotype-specific vulnerability.

The utility of this model extends to translational applications. Treatment with the IGFBP inhibitor NBI 31772 reduced astrogliosis, microglial activation, and neuronal loss, particularly in APOE3 Masteroids, but paradoxically increased tau phosphorylation and misfolding. These results mirror the differential efficacy and resistance observed among specific genotypic populations in human clinical trials, underscoring the predictive value of the Masteroid system for evaluating genotype-dependent therapeutic responses. By capturing both the protective and deleterious effects of interventions, our model provides a platform for optimizing drug candidates before clinical testing.

Overall, the Masteroid system demonstrates remarkable versatility, allowing for the study of individual AD components, genotype-specific effects, and complex interactions between Aβ, tau, and APOE4. It faithfully recapitulates key features of human AD, including amyloid plaque formation, tau pathology, neuroinflammation, and neuronal loss, while providing a rapid and reproducible platform for mechanistic studies and therapeutic testing. By bridging the gap between *in vitro* and *in vivo* models, Masteroids hold significant promise for advancing our understanding of AD pathogenesis and for accelerating the development of genotype-tailored therapies.

## Limitations of the study

While Masteroids capture key neuronal and glial components and recapitulate AD pathology, they lack full cellular diversity, vascularization, and long-range brain connectivity. Electrophysiological and network properties may not fully reflect the adult human brain, and the accelerated *in vitro* disease timeline may not model slow, chronic processes. Systemic factors and peripheral immune interactions are absent, and therapeutic responses may not fully predict clinical outcomes. Despite these limitations, Masteroids provide a reproducible, manipulable human-relevant platform for studying genotype-specific mechanisms and testing candidate interventions.

## Resource availability

### Lead contact

Further information and requests for resources and reagents should be directed to and will be fulfilled by the lead contact, Lulu Jiang.

### Materials availability

This study did not generate new unique reagents.

### Data and code availability

Single cell RNA-seq data generated in this study will be deposited in the NCBI Gene Expression Omnibus (GEO) and made publicly available upon publication.

This paper used existing R and STRING code packages and does not report original code. Any additional information required to reanalyze the data reported in this paper is available from the lead contact upon request.

## Acknowledgments

Post-mortem human brain tissues were provided by the Emory University Goizueta Alzheimer’s Disease Research Center (P30 AG066511). We thank Dr. Rakez Kayed and his lab for their generous support in providing the TOMA2 antibody. We also acknowledge the late Dr. Peter Davies (Northwell/Hofstra) and the Feinstein Institutes for Medical Research for providing the CP13 and MC1 antibodies. Electron Microscopy sample preparation was performed using the Leica UC7 at the Molecular Electron Microscopy Core, supported by the University of Virginia School of Medicine (RRID: SCR_019031). The RNA sequencing was conducted at the Genome Analysis and Bioinformatics Core at UVA (RRID:SCR_018883). We are grateful for the following funding support to L.J.: UVA Provost Award, UVA Health System, Virginia Alzheimer’s disease Center 2023 ‘Pitch and Catch’ funding, UVA Brain Institute Transformative Neuroscience Pilot Grant, Jim and Bruce Eck Fund, Strang Neuroscience Research Award, NIH/NIA (R01AG091577), Commonwealth Health Research Board Award, Cure Alzheimer’s Fund.

Data collection and sharing for *Figure 1* of this project was funded by the Alzheimer’s Disease Neuroimaging Initiative (ADNI) (National Institutes of Health Grant U01 AG024904) and DOD ADNI (Department of Defense award number W81XWH-12-2-0012). ADNI is funded by the National Institute on Aging, the National Institute of Biomedical Imaging and Bioengineering, and through generous contributions from the following: AbbVie, Alzheimer’s Association; Alzheimer’s Drug Discovery Foundation; Araclon Biotech; BioClinica, Inc.; Biogen; Bristol-Myers Squibb Company; CereSpir, Inc.; Cogstate; Eisai Inc.; Elan Pharmaceuticals, Inc.; Eli Lilly and Company; EuroImmun; F. Hoffmann-La Roche Ltd and its affiliated company Genentech, Inc.; Fujirebio; GE Healthcare; IXICO Ltd.; Janssen Alzheimer Immunotherapy Research & Development, LLC.; Johnson & Johnson Pharmaceutical Research & Development LLC.; Lumosity; Lundbeck; Merck & Co., Inc.; Meso Scale Diagnostics, LLC.; NeuroRx Research; Neurotrack Technologies; Novartis Pharmaceuticals Corporation; Pfizer Inc.; Piramal Imaging; Servier; Takeda Pharmaceutical Company; and Transition Therapeutics. The Canadian Institutes of Health Research is providing funds to support ADNI clinical sites in Canada. Private sector contributions are facilitated by the Foundation for the National Institutes of Health (www.fnih.org). The grantee organization is the Northern California Institute for Research and Education, and the study is coordinated by the Alzheimer’s Therapeutic Research Institute at the University of Southern California. ADNI data are disseminated by the Laboratory for Neuro Imaging at the University of Southern California.

## Author contributions

Conceptualization: L.J.

Investigation: E.S., K.Q., R.R., A.Z., A.E., L.J., L.S., S.L., H.S., L.I., W.G., K.S.

Visualization: E.S., K.Q., R.R., L.S., H.S., A.Z., A.E., L.J.

Data analysis: E.S., K.Q., L.S., S.L., A.Z.

Writing & Editing: E.S., L.J., K.Q., R.R., L.S., S.L., H.S., A.T., A.Z., A.E., J.K., J.S.

Supervision: L.J.

## Declaration of interests

L.J. and E.S. have filed a patent application related to the technology described in this manuscript. The authors declare no other competing interests.

## STAR METHODS

### KEY RESOURCES TABLE in the supplemental material

#### LEAD CONTACT AND MATERIALS AVAILABILITY

Further information and requests for resources should be directed to and will be fulfilled by the Lead Contact, Lulu Jiang.

### EXPERIMENTAL MODEL AND SUBJECT DETAILS

#### Cell Culture

##### Cell maintenance

All cell cultures were maintained at 37°C with 5% CO2. All cell counts were performed in duplicates using the C100 Automated Cell Counter (RWD) with trypan blue.

##### Human induced pluripotent stem cells (iPSC)

Human iPSCs were obtained from Jackson Laboratory (JIPSC001150, JIPSC001162) and maintained as a monolayer culture in StemFlex™ medium (Thermo Fisher Scientific, cat# A3349401) with 1% Penicillin-Streptomycin (Thermo Fisher Scientific, cat# 15070063) on Corning® Matrigel® hESC-qualified Matrix (Corning, cat# 354277) coated tissue culture plates. 1X RevitaCell™ Supplement (Thermo Fisher Scientific, cat# A2644501) was added to the medium when thawing. iPSCs were passaged at 80% confluency by Accutase™ (Stemcell Technologies, cat# 07920) dissociation as necessary. A full media change was performed every other day. iPSCs were cryopreserved in CryoStor® CS10 (Stemcell Technologies, cat# 07959). All iPSCs used for experimentation were maintained at passage <10.

##### iPSC neuronal progenitor cell (hiNPC)

Human iPSCs were passaged and plated at 50,000 cells/cm^2^ in STEMdiff™ SMADi Neural Induction medium (Stemcell Technologies, cat# 08581) with 10µM ROCK inhibitor Y-27632 (Stemcell Technologies, cat# 72304) and 1% Penicillin-Streptomycin on Corning® Matrigel® hESC-qualified Matrix coated tissue culture plates. hiNPCs were maintained in monoculture and passaged at 90% confluency by Accutase™ dissociation as necessary. A full media change was performed every other day. hiNPCs were cryopreserved STEMdiff™ Neural Progenitor Freezing Medium (Stemcell Technologies, cat# 05838).

##### iPSC neuronal cell (hiNeuron)

After 18 days in STEMdiff™ SMADi Neural Induction medium, hiNPCs were passaged and plated at 50,000 cells/cm^2^ in STEMdiff™ Forebrain Neuron Differentiation Medium (Stemcell Technologies, cat# 08600) with 10µM ROCK inhibitor Y-27632 and 1% Penicillin-Streptomycin on Corning® Matrigel® hESC-qualified Matrix coated tissue culture plates and transduced with a NEUROG2 lentivirus (GeneCopoeia, cat# CLP-HPRM50462-LvPF02-A00) at MOI 2 to induce iPSC-derived neuronal cells (hiNeurons). After 48 hours, the cells were again transduced using the same NEUROG2 lentivirus to ensure complete transduction. A full media change was performed 24 hours after each transduction.

##### iPSC astrocytic cell (hiAstrocyte)

iPSC-derived astrocyte progenitor cells (hiAPCs) were differentiated from hiNPCs. After 19 days in STEMdiff™ SMADi Neural Induction Medium, NPCs were passaged and plated at 50,000 cells/cm2 in STEMdiff™ Astrocyte Differentiation Medium (Stemcell Technologies, cat# 100-0013) with 1% Penicillin-Streptomycin on Corning® Matrigel® hESC-qualified Matrix coated tissue culture plates. A full media change was performed daily for 5 days, then every other day for 15 days, passaging as needed with Accutase™ dissociation. The differentiated hiAPCs were cryopreserved in CryoStor® CS10. At the time of experimentation, hiAPCs were thawed in STEMdiff™ Astrocyte Maturation Medium (Stemcell Technologies, cat# 100-0016) with 1% Penicillin-Streptomycin on Corning® Matrigel® hESC-qualified Matrix coated tissue culture plates. A full media change was performed every other day for 5 days with one passage by Accutase™ at 90% confluency as necessary.

##### iPSC hematopoietic progenitor cell (hiHPC)

iPSC derived hematopoietic progenitor cells (hiHPCs) were differentiated from iPSCs. Human iPSCs were plated in StemFlex medium with 1% Penicillin-Streptomycin as small aggregates on Corning® Matrigel® hESC-qualified Matrix coated tissue culture plates. The next day, the StemFlex medium was removed and replaced with STEMdiff™ Hematopoietic Basal Medium containing Supplement A (Stemcell Technologies, cat# 05310) and 1% Penicillin-Streptomycin. After two days, a half media change was performed. The following day, considered day 3, the medium was changed to STEMdiff™ Hematopoietic Basal Medium containing Supplement B and 1% Penicillin-Streptomycin. A half media change was performed on days 5, 7, and 10. hiHPCs were harvested at day 12 and either directly differentiated into microglia or cryopreserved in CryoStor® CS10.

##### iPSC microglial cell (hiMicroglia)

iPSC derived microglia (hiMicroglia) were derived from hiHPCs. hiHPCs were plated in STEMdiff™ Microglia Differentiation Medium (Stemcell Technologies, cat# 100-0019) and 1% Penicillin-Streptomycin on Corning® Matrigel® hESC-qualified Matrix coated tissue culture plates. 1mL of medium was added every 2-3 days. After 12 days, the cell suspension was collected and replated in the same medium. After 24 days in differentiation medium, the hiMicroglia were cryopreserved in CryoStor® CS10. Four days prior to experimentation, hiMicroglia were thawed in STEMdiff™ Microglia Maturation Medium (Stemcell Technologies, cat# 100-0020) with 1% Penicillin-Streptomycin on Corning® Matrigel® hESC-qualified Matrix coated tissue culture plates. 1mL of medium was added every other day.

##### Masterioid generation and maintenance

A single cell suspension of hiNeurons and hiAstrocytes was prepared by Accutase™ dissociation and washed once with DMEM/F12 (Stemcell Technologies cat# 36254) to remove debris, as described before^37^. A single cell suspension of hiMicroglia was prepared by collecting the floating cells and washing once with DMEM/F12. hiNeurons, hiAstrocytes, and hiMicroglia were combined in a 45:45:10 ratio in Masteroid medium ((DMEM/F12 (Stemcell Technologies, cat# 36254), 1% Glutamax (Thermo Fisher Scientific, cat# 35050061), 1% sodium pyruvate (Thermo Fisher Scientific, cat# 11360070), 1% N-2 Supplement (Thermo Fisher Scientific, cat# 17502-048), 1% B-27 Supplement (Thermo Fisher Scientific, 17504044), 10µM Y-27632,1% Penicillin-Streptomycin, 1mg/mL Heparin (Sigma-Aldrich, cat# H3149-250KU), and 10% STEMdiff™ Microglia Maturation Medium), passed through a 40µm cell strainer, and plated at 2,000 cells/microwell in AggreWell™800 microwells (Stemcell Technologies, cat# 34815) coated in Anti-Adherence Rinsing Solution (Stemcell Technologies, cat# 07010). The Aggrewell™ plate was immediately centrifuged at 100×g for 3 minutes to capture the cells in the microwells and incubated for 24 hours. A half-media change was performed at 24 hours and then every other day for 1 week. At 1 week when the spheroids displayed a smooth, bright edge under the cell culture microscope, cultures were transferred to ultra-low-attachment round-bottom 96-well plates (Thermo Fisher Scientific, cat# 174929) and maintained in 200µL Masteroid media rotating at 65rpm.

For the generation of Masteroids with AD oTau seeding, hiNeurons were exposed to 40ng/mL oTau fraction, extracted from post-mortem AD brain tissues or age-matched normal control, by direct admission into cell culture media for 24 hours prior to their incorporation into Masteroid. The concentration of oTau fraction to be added into the iNeurons was determined by dose-response test. For the Masteroids with Aβ oligomers conditions, Masteroids were exposed to 0.5µM Aꞵ oligomers or vehicle control (DMSO) after 1 week when the Masteroids were transferred to 96-well plates (Anaspec, cat# AS-64129-05). The dose of Aβ oligomers were determined by a dose-response test in hiNeurons and hiAstrocyte.

For NBI 31772 treatment, Masteroids were exposed to 0.5µM of NBI 31772 (MedChemExpress, cat# HY-110135) or vehicle control (DMSO) at 5 days before collection. Half media change was performed daily with 1µM NBI 31772 to maintain the 0.5µM concentration.

##### 2D neuron and glial culture treatment

At the time point of creating the Masteroids, individual cultures of hiNeurons, hiAstrocytes, and hiMicroglia were passaged and plated in Masteroid media onto Corning® Matrigel® hESC-qualified Matrix coated tissue culture 8 well chambers. Half media changes were performed every other day.

#### Oligomer processing

##### Generation of S1p oTau fraction

The anonymous post-mortem human brain tissues used in this study were sourced from the brain bank of the Emory Goizueta Alzheimer’s Disease Research Center (ADRC, funded by NIH P30AG066511). All fresh frozen tissue samples (8 age-matched control and 8 AD) were de-identified and obtained from the human prefrontal cortex tissues (Brodmann Area 10) within the same patient cohort. These brain regions were selected due to their consistent availability in the brain bank and previous research indicating high positive correlations between altered gene expression and disease severity measures, regardless of the classification scheme for disease severity^113^. The generation of S1p oTau fractions was consistent with previous publications^68,70^. In brief, frozen sections were weighed (100-250mg) and put into polycarbonate thick wall 1.5mL tubes (Beckman Coulter, cat# 357448). A 10x volume of homogenization buffer, Hsaio TBS buffer (50mM Tris, pH 8.0, 274mM NaCl, 5mM KCl) supplemented with cOmplete™ Protease Inhibitor Cocktail (Roche, cat# 05892791001) and the phosphatase inhibitor cocktail PhosSTOP™ (Roche, cat# 04906837001) was used to homogenize brain tissue. The homogenate was centrifuged at 29,800 x g for 20 minutes at 4°C. The supernatant was then centrifuged a second time at 150,000 x g at 4°C for 40 minutes. The TBS-extractable pellet (S1p) fraction was resuspended with 50uL TE buffer (10mM Tris, pH 8.0, 1mM EDTA, 5mM KCl) supplemented with cOmplete™ Protease Inhibitor Cocktail and PhosSTOP™ relative to the starting weight of the tissue, homogenized, aliquoted, and frozen at −80°C. Tau oligomerization was confirmed through running a western blot and visualizing total, misfolded, and oligomerized tau with Anti-Tau Antibody clone Tau-5 (Millipore Sigma, cat# MAB361) and Phospho-Tau (Thr217) Polyclonal Antibody (Thermo Fisher Scientific, cat# 44-744).

The human brain tissue samples used in this study were all de-identified. All studies included both sexes, and results were integrated by covariate analysis, as described below.

| **Primary Neuropathologic Diagnosis** | **Braak Stage** | **Age at Onset** | **Age at Death/Bx** | **Duration (years)** | **APOE** | **Race** | **Sex** |
| --- | --- | --- | --- | --- | --- | --- | --- |
| Alzheimer's disease (AD) | VI | 49 | 59 | 10 | E3/3 | w | m |
| AD | VI | 59 | 69 | 10 | E3/4 | w | m |
| AD | VI | 53 | 71 | 18 | E3/4 | w | m |
| AD | VI | 65 | 80 | 15 | E3/4 | b | f |
| AD | V | 77 | 83 | 6 | NA | w | f |
| AD | V | 86 | 92 | 6 | E3/4 | w | f |
| AD | V | 82 | 92 | 10 | E3/3 | w | m |
| AD | VI | 80 | 93 | 13 | E3/4 | w | f |
| Normal Control (NC) | I | NA | 59 | NA | E2/3 | b | m |
| NC | I | NA | 70 | NA | E3/3 | b | m |
| NC | I | NA | 72 | NA | E3/3 | w | m |
| NC | II | NA | 78 | NA | E3/3 | w | f |
| NC | IV | NA | 84 | NA | E2/3 | b | f |
| NC | III | NA | 91 | NA | E3/3 | w | f |
| NC | III | NA | 92 | NA | E3/3 | w | f |
| NC | II | NA | 94 | NA | E3/3 | w | m |

##### Preparation of Aꞵ oligomers

0.5mg of human beta-Amyloid 1-42 (Anaspec, cat# AS-64129-05) was dissolved in 50µL of filtered DMSO then transferred to a 1.5-mL Protein LoBind Eppendorf tube (Eppendorf, cat# 022-43-108-1). The dissolved Aβ was sonicated at a high power (550 Sonic Dismembrator, machine output power 550W, output frequency 20KHz) on ice for 2 minutes then resuspended with 60µL of neurobasal medium (Thermo Fisher Scientific, cat# 21103049) for a final concentration of 1mM. The resuspension was rotated at 4°C for 24 hours to promote oligomerization. Oligomerization was confirmed through running a native-page gel and visualizing with Purified anti-β-Amyloid, 1-16 Antibody (BioLegend, cat# 803001). Aꞵ oligomers were aliquoted and stored at −80°C.

#### Sample collection

##### Masteroid fixation for immunofluorescence

At the time of collection, Masteroids were transferred to a 1.5mL Protein LoBind Eppendorf tube (Eppendorf, cat# 022-43-108-1) and allowed to settle. The supernatant was discarded, and the Masteroids were fixed in 4% PFA in 1x PBS for 15 minutes. After fixation, Masteroids were washed 3 times for 10 minutes each in 1x PBS. Samples were stored in 1x PBS at 4°C.

##### Masteroid fixation for electron microscopy

At the time of collection, Masteriods were transferred to a 1.5mL tube and allowed to settle. The supernatant was discarded, and the Masteroids were fixed with 4% PFA and 2.5% glutaraldehyde (Electron microscopy sciences, cat# 16316) in 1x PBS for 20 minutes. After fixation, Masteroids were washed 2 times for 10 minutes each in 1x PBS. Samples were stored in 4% PFA in 1x PBS at 4°C.

##### 2D individual culture fixation for immunofluorescence

After 1 week in culture, the individual cell types were washed once with 1x PBS then fixed in 4% PFA in 1x PBS for 15 minutes. After fixation, chambers were washed 3 times for 10 minutes each in 1x PBS. Samples were stored in 1x PBS at 4°C.

##### Protein lysate

At the time of collection, 16 pooled Masteroids per condition were transferred to a 1.5mL Protein LoBind Eppendorf tube and allowed to settle. The supernatant was discarded, and the Masteroids were washed once with PBS. The PBS was discarded, and the samples were flash-frozen and stored at −80°C. Samples were subsequently lysed by mechanical homogenization in 40µL of Hsiao TBS buffer (50mM Tris, pH 8.0, 274mM NaCl, 5mM KCl) supplemented with cOmplete™ Protease Inhibitor Cocktail (Roche, cat# 05892791001) and the phosphatase inhibitor cocktail PhosSTOP™ (Roche, cat# 04906837001). 200U of Benzonase (Sigma-Aldrich, cat# E1014-25KU) with 1mM MgCl2 was added for removal of nucleic acids. Lysate concentration was quantified by the Pierce BCA assay according to the manufacturer’s protocol (Thermo Fisher Scientific, cat# 23225).

##### 14µm thin sectioning of Masteroids

Masteroids were pooled per condition and fixed in a freshly prepared solution of 4% paraformaldehyde and 0.1% glutaraldehyde in PBS for 5 minutes at room temperature. After fixation, samples were washed three times in PBS for 10 minutes each. A thin layer of a 3% agar solution, maintained at 55°C, was poured into a 35mm petri dish. Masteroids were carefully transferred to the petri dish and positioned on the same layer no more than 1cm apart, and additional 3% agar solution was used to fully embed the Masteroids then cooled for 15 minutes to solidify. The block containing the Masteroids was separated using a razor blade, then carefully extracted with a spatula. To prevent freezing artifacts during cryosectioning, Masteroids were equilibrated in 30% sucrose for 24 hours at 4°C. Embedded samples were sectioned at 14µm thickness using a low-profile blade in a cryochamber at −30°C and collected onto positively charged glass slides. Slides with Masteroid sections were stored at 4°C.

##### Conditioned media collection

60µL of conditioned cell culture media from replicate Masteroid wells were collected in flat-bottomed 96-well plates and frozen at −20°C.

#### Sample processing

##### Immunofluorescence labeling

Selected Masteroids from each condition were washed in 150µL PBS for 10 minutes in a U-bottom 96-well plate and then permeabilized in 150µL PBS/0.25% Triton X-100 (PBST) for 30 minutes. The Masteroids were blocked with 150µL PBST with 5% Bovine Serum Albumin (BSA) and 5% Normal Goat Serum (NGS) for 1 hour at room temperature (RT). After blocking, the Masteroids were incubated in primary antibodies diluted in 5% BSA in PBST overnight at 4°C.

Plates were wrapped with parafilm and photobleached by an LED lamp. The following day, the Masteroids were washed 3 times with PBST for 10 minutes each before being incubated in secondary antibodies diluted (1:1000 of Alexa-conjugated antibodies made from goat purchased from Thermo Fisher Scientific) in PBST with 5% BSA for two hours. The Masteroids were protected from light and washed 3 times with PBST for 10 minutes each followed by one 10-minute wash with PBS. For Masteroids stained with DAPI Nucleic Acid Stain (Thermo Fisher Scientific, cat# 62248, 1:5,000 of 1mg/mL stock) was diluted in PBST and incubated with asteroids for 15 minutes, followed by 2× washes with PBST then 1× with PBS, 10 minutes each. Masteroids were then mounted onto microscope glass slides in Prolong™ Gold Antifade Mountant (Thermo Fisher Scientific, cat# P36934) and protected from light. All washing and incubated steps were done with agitation. Primary antibodies used for Masteroid immunofluorescent labeling were as follows: Tuj1/Anti-Beta-Tubulin 3 Antibody (chicken, Aves Labs, cat# TUJ, 1:500), GFAP Monoclonal Antibody (rat, Thermo Fisher Scientific, cat# 13-0300, 1:500), Anti Iba1 (rabbit, FUJIFILM Wako Pure Chemical Corporation, cat# 019-19741, 1:300), Anti Iba1 (goat, FUJIFILM Wako Pure Chemical Corporation, cat# 011-27991, 1:300), Monoclonal anti-TOMA2 (mouse, provided by Dr. Rakez Kayed, 1:500), Myelin Basic Protein (rabbit, Cell Signaling Technology, cat# 78896, 1:500), 6E10/Purified anti-β-Amyloid, 1-16 Antibody (mouse, BioLegend, cat# 803001, 1:500), CD11b Antibody, clone 5C6 (rat, Bio-Rad, cat# MCA711G, 1:300), Anti-P2Y12 Antibody (rabbit, Abcam, cat# ab300141, 1:500), and Anti-Microtubule-Associated Protein Antibody (chicken, Antibodies Incorporated, cat# MAP, 1:500). Secondary antibodies used for Masteroid immunofluorescent labeling were as follows: Goat anti-Rat IgG (H+L), Alexa Fluor 405 (Thermo Fisher Scientific, cat# A48261), Goat anti-Rat IgG (H+L), Alexa Fluor 488 (Thermo Fisher Scientific, cat# A11006), Goat anti-Rabbit IgG (H+L), Alexa Fluor 488 (Thermo Fisher Scientific, cat# A11008), Goat anti-Rabbit IgG (H+L), Alexa Fluor 594 (Thermo Fisher Scientific, cat# A11012), Goat anti-Mouse IgG (H+L), Alexa Fluor 405 (Thermo Fisher Scientific, cat# A48255), Goat anti-Mouse IgG (H+L), Alexa Fluor 488 (Thermo Fisher Scientific, cat# A28175), Donkey anti-Goat IgG (H+L), Alexa Fluor 647 (Thermo Fisher Scientific, cat# A21447), and Goat anti-Chicken IgG (H+L), Alexa Fluor 647 (Thermo Fisher Scientific, cat# A32933). More details are available from the Key Research Table in the supplemental materials.

##### Immunofluorescence staining of 14µm thin sections of Masteroids

Masteroids were sliced and collected onto slides as described in the “14µm thin sectioning of Masteroids” protocol. Sections were outlined with a hydrophobic barrier using a PAP pen and air-dried before staining. Immunofluorescence staining was performed as described in the “Immunofluorescence Staining” protocol, with the modification of using 0.1% Triton X-100 in PBS instead of 0.25%.

*Immunohistochemistry single stain with DAB (3,3’-diaminobenzidine) of 14µm thin sections of Masteroids*

Masteroids were sliced and collected onto slides as described in the “14µm thin sectioning of Masteroids” protocol. All subsequent steps were performed at room temperature unless otherwise specified. Sections were outlined with a hydrophobic barrier using a PAP pen and air-dried before staining. Sections were washed in PBS for 10 minutes then treated with 1% H_2_O_2_ diluted in dH2O for 15 minutes. Sections were rinsed twice in PBS for 15 minutes each, then blocked with 0.4% Triton X-100 with 1% BSA and 4% NGS in PBS for 30 minutes. After blocking, sections were incubated overnight in 1° antibody diluted in DAKO antibody diluent (5% BSA in 0.25% TritonX-100/PBS) at 4°C with photobleaching. The following day, sections were rinsed twice in PBS for 15 minutes then incubated for one hour in BTA solution (0.44% Biotinylated goat anti-rabbit / mouse (depending on 1° antibody) IgG diluted in 0.3% Triton X-100/PBS). Sections were rinsed twice in PBS then treated with Avidin Biotin Complex (ABC) (0.88% Avidin, 0.88% Biotin diluted in 0.3% Triton X-100/PBS, prepared 30 minutes prior to use; Vector Laboratories, cat# PK-6101). After ABC incubation, sections were washed three times in PBS for 10 minutes each. DAB solution was prepared by adding 1 tablet DAB (Millipore Sigma, cat# D4293-5SET) and 1 tablet Urea (Sigma-Aldrich, cat# D4193) to 5mL dH_2_O and vortexed until fully dissolved. DAB solution was added to sections individually for one to three minutes until section turned medium brown in color, then were immediately placed in PBS. Sections were washed twice with PBS for five minutes each, then dried overnight at 37°C. The following day, sections were dehydrated by washing in 70% EtOH, 85% EtOH, 95% EtOH, and 100% EtOH for two minutes each followed by washing in 100% EtOH for five minutes. Slides were then cleared by washing twice in Xylene for five minutes followed by immediately dropping 50µL of Permount Mounting Medium (Thermo Fisher Scientific, cat# SP15-500) and covering with glass coverslips. Slides were dried overnight in the fume hood and stored at 4°C for long term storage. All washing and incubated steps were done with agitation. Primary antibodies used for Masteroid DAB labeling were as follows: MC1 (mouse, Dr. Peter Davies and the Feinstein Institutes for Medical Research, 1:300) and Phospho-Tau (Thr217) Polyclonal Antibody (rabbit, Thermo Fisher Scientific, cat # 44-744, 1:500). Secondary antibodies used for Masteroid DAB labeling were as follows: Goat anti-Rabbit IgG Antibody (H+L), Biotinylated (Vector Laboratories, cat# BA-1000) and Goat anti-Mouse IgG Antibody (H+L), Biotinylated (Vector Laboratories, cat# BA-9200). More details are available from the Key Research Table in the supplemental materials.

##### Electron microscopy

The protocol for sample preparation is consistent with previous publications^114^. In brief, Resin embedding for electron microscopy followed standard protocols. Masteroids fixed with 4% PFA and 2.5% glutaraldehyde (EMS, cat# 16316) were transferred to Beem Capsules (EMS, cat# 70010-B) in 0.1M phosphate buffer (PB) and allowed to settle. All subsequent steps were carried out in the same Beem Capsules. About half of the buffer was removed and replaced with 1% osmium tetroxide in 0.1M PB, then once the Masteroids were visible, all the buffer was removed and replaced with the 1% osmium tetroxide solution for one hour. The Masteroids were then transferred into 50% EM grade ethanol (EMS, cat# 15055) for three minutes and counterstained with filtered 4% uranyl acetate in 70% ethanol overnight, in the refrigerator. The sections were dehydrated through a series of ethanol and EM grade acetone (EMS, cat# 10015) solutions and infiltrated with a 1:1 ratio of EPON EMbed 812 resin (EMS, cat# 14121) and acetone overnight. They were then transferred to 100% EPON resin overnight. Capsules were then placed in a 60°C oven for 48 hours until hardened, the resin block is removed from the capsules, the block face was trimmed to 1×2mm that contains multiple Masteroids, and then cut into ultrathin (60-70nm) sections using an ultramicrotome (Leica, Ultracut UCT7). The ultrathin sections were collected on 200 mesh copper grids (EMS, cat# Cu G200TT-Cu) and slot copper grids (EMS, cat# FCF-2010-Cu-50) in ultrapure water and allowed to dry before imaging.

##### Immunoblot/ Western blot

10µg of Masteroid lysate was separated by denaturing gel electrophoresis (SDS-page) and transferred onto a nitrocellulose membrane. The membrane was blocked for an hour in 5% non-fat milk in .01% Tween-20 in 1x PBS (PBST), and incubated in primary antibody (dilution antibody dependent) in 5% BSA PBST overnight at 4°C. The following day, the membrane was washed with PBST for 3 times for 10 minutes each then incubated with secondary antibody (1:5000) conjugated with horseradish peroxidase (Thermo Fisher Scientific, cat# G-21040; cat# A16054; cat# G-21234; cat# A16054) in 5% BSA PBST for 2 hours at RT. The membrane was washed with PBST 3 times for 10 minutes before briefly being washed with PBS prior to visualization. Antibody staining was visualized using SuperSignal™ West Pico PLUS Chemiluminescent Substrate (Thermo Fisher Scientific, cat# 34578) and imaged by ChemiDoc Imaging System (Bio-Rad). For reprobing, membranes were incubated in Restore™ PLUS Western Blot Stripping Buffer (Thermo Fisher Scientific, cat# 46430) for 15 minutes at RT. Band intensities were quantified using ImageJ. Protein abundance was normalized to total Beta Actin with background subtraction. Primary antibodies used for immunoblotting were as follows: IBA1 Polyclonal Antibody (rabbit, Proteintech, cat# 10904-1-AP, 1:1000), GFAP Monoclonal Antibody (rat, Thermo Fisher Scientific, cat# 13-0300, 1:1000), 6E10/Purified anti-β-Amyloid, 1-16 Antibody (mouse, BioLegend, cat# SIG-39320, 1:1000), Beta Actin Monoclonal Antibody (mouse, Proteintech, cat# 66009-1-Ig, 1:2000), NeuN Polyclonal Antibody (rabbit, Thermo Fisher Scientific, cat# PA5-78499, 1:1000), and CP13 (mouse, Dr. Peter Davies and the Feinstein Institutes for Medical Research, 1:1000). Secondary antibodies used for immunoblotting were as follows: Goat anti-Rabbit IgG (H+L), HRP (Thermo Fisher Scientific, cat# G21234, 1:2000), Goat anti-Mouse IgG (H+L), HRP (Thermo Fisher Scientific, cat# G21040, 1:2000), and Goat anti-Rat IgG (H+L), HRP (Thermo Fisher Scientific, cat# A18865, 1:2000). More details are available from the Key Research Table in the supplemental materials.

##### LDH cytotoxicity assay

The CyQUANT™ LDH Cytotoxicity Assay was performed as per manufacturer’s instructions using 50µL conditioned media replicates to measure lactate dehydrogenase (LDH) release (Thermo Fisher Scientific, cat# C20301). The assay 490nm and 680nm absorbance readings were taken on an Epoch microplate spectrophotometer (BioTek).

##### Single-cell RNA-sequencing sample preparation

The protocol for Single-cell RNA-sequencing sample preparation was consistent from our previous publication^37^. In brief, for each condition, 30 Masteroids were pooled in a 1.5-mL Protein LoBind Eppendorf tube and allowed to settle. The supernatant was discarded, and a single-cell suspension was produced by incubation in 500µL of digestion buffer (Accutase™ with 80 U/mL Protector RNase Inhibitor (Sigma Aldrich, cat# 3335399001)) for 1 hour at 37°C with gentle pipette mixing every 10 minutes. At the end of the incubation the single-cell suspension was washed with 500μL wash buffer (0.02% BSA in 1X PBS with 80 U/mL Protector RNase Inhibitor) and passed through a 20μM filter (Miltenyi Biotec, cat# 130-101-812) to a fresh 2-mL Protein LoBind Eppendorf. After another centrifugation, the supernatant was discarded, and the single-cell pellet gently resuspended in 50μL of wash buffer. Cells were counted in triplicate on the Nexcelom Cellometer Auto 2000 with trypan blue and processed through the single-cell RNA-sequencing pipeline from 10X Genomics, 3’ Version 3 (10X Genomic Chromium).

##### Single-cell RNA-sequencing library preparation and sequencing

The single-cell RNA library preparation and sequencing was done by the University of Virginia School of Medicine’s Genome Analysis and Technology Core (GATC; RRID:SCR_018883). cDNA library creation: RNA samples were processed for single-cell sequencing according to manufacturer’s instructions using the kit, according to validated standard operating procedures established by the GATC. cDNA quality assessment: cDNA quality was assessed by the MiSeq Reagent Nano Kit v2 (Illumina, cat# MS-102-2001). Single-cell libraries were sequenced using a p4-100 kit (Chromium Next GEM Single Cell 3’ GEM, Library & Gel Bead Kit v3.1, 16 rxns, Illumina, cat# PN-1000121) on the NextSeq2000 Sequencing System (Illumina).

##### Whole-cell recording of Masteroids

Neuronal activity in whole neuron-glial brain assembloid was recorded using a whole-cell patch clamp technique as described previously^115^. Whole-cell patch-clamp recordings were performed under infrared differential interference contrast microscopy (Olympus); a 40× water-immersion objective was used to visually identify neurons in the brain assembloid. The Masteroids were continuously perfused with ACSF solution that was saturated with 95% O2 and 5% CO2 at room temperature. Patch electrodes (final resistances, 3–5 MΩ) were pulled from borosilicate glass (Sutter Instruments, Novato, CA, USA) on a horizontal Flaming-Brown microelectrode puller (Model P-97, Sutter Instruments). For voltage-clamp recordings, the electrode tips were filled with a filtered internal recording solution that consisted of the following components (in mM): 100 Cs-gluconate, 10 Hepes, 0.6 EGTA, 4 MgATP and 0.3 NaGTP; 5 QX-314 the pH was 7.3 (with CsOH), and the osmolarity was 310 mosmol l−1. For current-clamp recording, the pipette solution contained the following (in mm): 135 potassium gluconate, 2.5 NaCl, 10 Hepes, 0.5 EGTA, 4.0 Mg-ATP, 0.4 Na-GTP, 0.1 CaCl2; the pH was 7.3, and the osmolarity was 310 mosmol l−1. Neurons were voltage clamped at −60mV using a PC-505B amplifier (Warner Instruments, Hamden, CT, USA). Electrode capacitance was electronically compensated. Access resistance was continuously monitored, and if the series resistance increased by 20% at any time, the recording was terminated. Currents were filtered at 2 kHz, digitized using a Digidata 1322 digitizer (Molecular Devices, Sunnyvale, CA, USA), and acquired using Clampex 10.2 software (Molecular Devices). Spontaneous excitatory postsynaptic currents (sEPSCs) were recorded from neurons after blocking the GABA_A_ receptors with the antagonist picrotoxin (50μm).

#### Quantification and statistical analysis

##### Fluorescent image analysis

Fluorescent images were captured by STELLARIS 5 Confocal Microscope (Leica Microsystems, Stellaris 5) and LP8 Confocal Microscope (Leica Microsystems, SP8) using LAS X Life Science Microscope Software Platform (Leica Microsystems). The staining intensities in immunofluorescence-labeled Masteroids were measured by ImageJ^116^.

##### Brightfield image analysis

Brightfield images (DAB) were captured by Keyence BZ-X810 Microscope (Keyence, BZ-X810) using BZ-X800 Viewer Software (Keyence). The staining intensities in DAB-labelled Masteroids were measured by ImageJ^116^.

##### Electron microscopy analysis

Electron microscopy (EM) images were captured on a JEOL 1010 EM with a 16-megapixel CCD camera (SIA) at magnifications ranging from 4kx to 20kx, yielding nm/pixel resolution. Adobe Photoshop was used to balance and annotate EM images. Cross-sections of dendrites were identified by the presence of microtubules and lack of neurotransmitter vesicles. Axon terminals may display presynaptic zones and contain vesicles. Glial and neuronal cells were identified based on established accepted criteria^61,65,117,118^.

##### Schematics

Schematics were created with BioRender.com

#### Data analysis

##### Statistical analysis

Statistical analyses and figures artwork were performed using GraphPad Prism version 10.00 for Windows. Pairwise data was compared using one-tailed Wilcoxon matched-pairs signed rank test. All group data are expressed as mean ± SEM. Column means were compared using one-way ANOVA with treatment as the independent variable. Group means were compared using two-way ANOVA using factors of genotype and fraction treatment, respectively. When ANOVA showed a significant difference, pairwise comparisons between group means were examined by Tukey’s, Dunnett, or uncorrected Fisher’s LSD multiple comparison test. Significance was defined when *p* < 0.05. Gene networks were visualized using the Cytoscape StringApp 2.2.0 STRING protein query through Cytoscape 3.10.3.

#### Single-cell RNA-sequencing analysis

##### CellRanger pipeline

CellRanger version 3.1.0 (10X Genomics) was used to combine and process the raw Illumina NextSeq 500 RNA and NOVAseq sequencing files. First, each sequencing library was demultiplexed by sample index to generate FASTQ files for paired-end reads using the CellRanger mkfastq pipeline. FASTQ files were then passed to the CellRanger count pipeline, which used STAR aligner version 2 to align reads to the human reference genome (GRCh38)^119^. Gene-cell barcode matrices of each sample were further processed in R version 4.0.3.

##### Seurat object filtration

Datasets were loaded into Seurat v5.0.3 for filtering, normalization, and data scaling^120^. Cells containing fewer than 2000 and greater than 20000 RNA counts, and greater than 25% mitochondrial RNA were filtered out using Seurat’s subset function. Datasets were processed using SCTransform and integrated using Seurat’s CCA integration^121^. Clusters were determined from the uniform manifold approximation and projection (UMAP) dimensional reduction graph via SLM algorithm using Seurat’s native FindClusters function at 1.5 resolution^122^.

##### Cluster cell-type identification

Identities of each cluster were labeled manually using the APOE3 NC condition through various methods, including comparing cluster marker genes to known human neuronal/glial cell types, scoring each cluster enrichment of marker genes from our previous single-cell organoid datasets using AddModuleScore^37,123^. Clusters otherwise not found in APOE3 NC were matched to the nearest “normal” cluster by querying the dataset containing all clusters to the annotated APOE3 NC-only dataset. Significantly upregulated genes in disease conditions were obtained using Wilcoxon ranked-sum testing.

##### Differential expression analysis

Differential expression analysis was performed within each cell type between different conditions using the Wilcoxon rank-sum test implemented in Seurat with a min.pct of 0.1, a logfc.threshold of 0.25, and a pseudocount of 1 (default). A multiple-comparison correction was performed using the Benjamin & Hochberg FDR method to produce an adjusted *p* value^124^. Differentially expressed genes were evaluated according to their log fold change (greater than log2(.25)) and adjusted *p* values (<0.05).

##### Functional enrichment analysis

Functional enrichment analysis of the significantly differentially expressed genes between different conditions was performed using the R implemented GProfiler version 0.2.3^125^. The enrichment analysis was run as an ordered query (ordered by log2FC) using an α threshold of 0.05 and using Benjamin & Hochberg FDR for multiple testing correction^124^. Only genes in the Seurat dataset were considered by using a custom domain scope. A custom source GMT, gp zSEF_sD9Q_d1M, was used. It includes all Hallmark gene sets, curated gene sets, and ontology gene sets from the Molecular Signatures Database (MsigDB) v7.2^126,127^.

##### Gene set enrichment analysis

To generate NES scores/assess pathway-level changes in gene expression across conditions, we performed GSEA within each cell type cluster and condition comparison^126^. DE analysis was conducted (as previously described) and all expressed genes were ranked by average log fold change between the two conditions. This ranked list was used as input to the fgseaMultiLevel function from the fgsea R package, which implements an adaptive multilevel approach for improved accuracy in estimating enrichment *p* values. We used the same custom MSigDB gene set collection as previously described, filtered to include only gene sets with at least one gene expressed in the given cluster. The analysis output included normalized enrichment scores and adjusted *p* values for each pathway.

##### CellChat analysis

To infer intercellular signaling networks, we used CellChat R package version 2.20 to analyze ligand-receptor mediated communication between cell types^57^. For each condition, we subsetted the Seurat object to include only cells from the selected condition and 5 major cell types. CellChat objects for each condition were then initialized using the RNA assay and grouped by cell type identity. The ‘secreted signaling’ subset of the human CellChatDB was used for this analysis. The general CellChat pipeline included: SubsetDatta to retain only signaling-related genes; identifyOverExpressedGenes and identifyOverExpressedInteractions to identify potential ligands and receptors; computeCommunProb to infer the probability of communication between cell types; computeCommunProbPathway to summarize interactions at the signaling pathway level; aggregateNet to construct a global signaling network (for each condition). We extracted both counts (total number of significant ligand-receptor interactions between each sender-receiver cell type pair) and weights (summed communication probabilities representing strength of signaling between cell pairs). Circle plots were generated for visualization of count and weights.

##### Cytoscape analysis

A STRING network of the significant (*p* value < 0.05) differentially expressed genes (DEGs) in a given cell cluster was created using the Cytoscape StringApp 2.2.0 STRING protein query through Cytoscape 3.10.3^128,129^. The network was then clustered by Markov clustering (MCL) with a granularity inflation of 4 to create gene nodes using clusterMaker2^130^. Functional enrichment was run on individual nodes using STRING enrichment to determine the enriched pathways, and nodes that were associated with similar pathways were grouped together. For visualization, manual annotations were assigned based on the most abundant or relevant pathways per node. The color gradient was applied using Continuous Mapping with the log2FC value of each gene. The most enriched 6-10 clusters were presented in the figure.

#### PET analysis

##### Tau PET analysis

T1-weighted (T1w) MRI and 18F-flortaucipir (AV-1451) PET images were downloaded from the ADNI repository (https://adni.loni.usc.edu). For T1w images, cross-sectional image processing was performed using FreeSurfer version 7^131–134^. Six five-minute frames were acquired 75 to 105 minutes post-injection. Individual frames were extracted from the image files and co-registered to the first extracted frame of the raw image file. The base frame and co-registered frames were recombined into a dynamic image set. The images were further processed using PETSurfer^131,132^, utilizing segmentation results from FreeSurfer. We applied partial volume correction (PVC) with a 6mm extent using geometric transfer matrix (GTM) methods. After segmentation and registration, PETSurfer calculates SUVR using the pons as the reference region. Average SUVR measures were calculated using the DKT^135,136^ and ASAG^137^ atlases for cortical and subcortical ROIs.

##### Amyloid PET analysis

Preprocessed ¹⁸F-florbetapir (AV-45) amyloid PET data were provided by the ADNI database. Specifically, we used amyloid positivity status based on the cutoff value of 1.11 to the global cortical AV-45 SUVR normalized by the whole cerebellum. These SUVR values were preprocessed using the ADNI UC Berkeley AV-45 pipeline^138^, where PET frames acquired 50–70 minutes post-injection were motion-corrected, averaged, co-registered to corresponding T1-weighted MRI scans, and spatially normalized.

##### Subject selection

Since ADNI is a longitudinal study that includes individuals who are cognitively unimpaired (CU), have mild cognitive impairment (MCI), or have dementia/Alzheimer’s disease (DEM), we selected subjects whose APOE genotype was APOE3/3, APOE3/4, or APOE4/4 and used their earliest available PET imaging data for analysis. Summarization of the distribution of subjects by APOE genotype and diagnostic group can be found in the Source Data.

##### ADNI Data Acquisition

PET imaging data used in the preparation of *Figure 1* of this article were obtained from the Alzheimer’s Disease Neuroimaging Initiative (ADNI) database (adni.loni.usc.edu). ADNI was launched in 2003 as a public-private partnership, led by Principal Investigator Michael W. Weiner, MD. The primary goal of ADNI has been to test whether serial magnetic resonance imaging (MRI), positron emission tomography (PET), other biological markers, and clinical and neuropsychological assessment can be combined to measure the progression of mild cognitive impairment (MCI) and early Alzheimer’s disease (AD).

## Abbreviations

AD: Alzheimer’s Disease
oTau: Oligomeric tau
NC: Normal Control
Aβ: Amyloid beta
AD oTau: Oligomeric tau derived from diseased AD brains
AD Aβ: Condition with both Aβ oligomers and AD oTau
UMAP: Uniform manifold approximation and projection
GSEA: Gene set enrichment analysis
CU: Cognitively unimpaired
MCI: Mild cognitive impairment
DEM: Demented
ADNI: Alzheimer’s Disease Neuroimaging Initiative

