## Supplemental Figures for "Modeling Alzheimer’s Disease with APOE4 Neuron-Glial Brain Assembloids Reveals IGFBPs as Therapeutic Targets"

*A list of authors and their affiliations appears at http://adni.loni.usc.edu/wp-content/uploads/how_to_apply/ADNI_Acknowledgement_List.pdf.

#### List of Supplemental Materials

Figure S1. APOE4 status significantly increases amyloid prevalence and tau burden by ADNI PET data, related to Figure 1.

Figure S2. Masteroids exhibit structural and biophysical properties reminiscent of the human brain, related to Figure 2.

Figure S3. Modification of Masteroid culture system creates AD model, related to Figure 3.

Figure S4. scRNA UMAPs and CellChat networks for all Masteroid conditions, related to Figures 2, 3, 4, 5, and 6.

Figure S5. APOE4 Masteroids exhibit expansive AD pathology, related to Figure 4.

Figure S6. Contribution of APOE4 in driving AD pathogenesis, related to Figures 5 and 6.

Figure S7. NBI 31772 impacts AD pathology in APOE4 Masteroids, related to Figure 7.

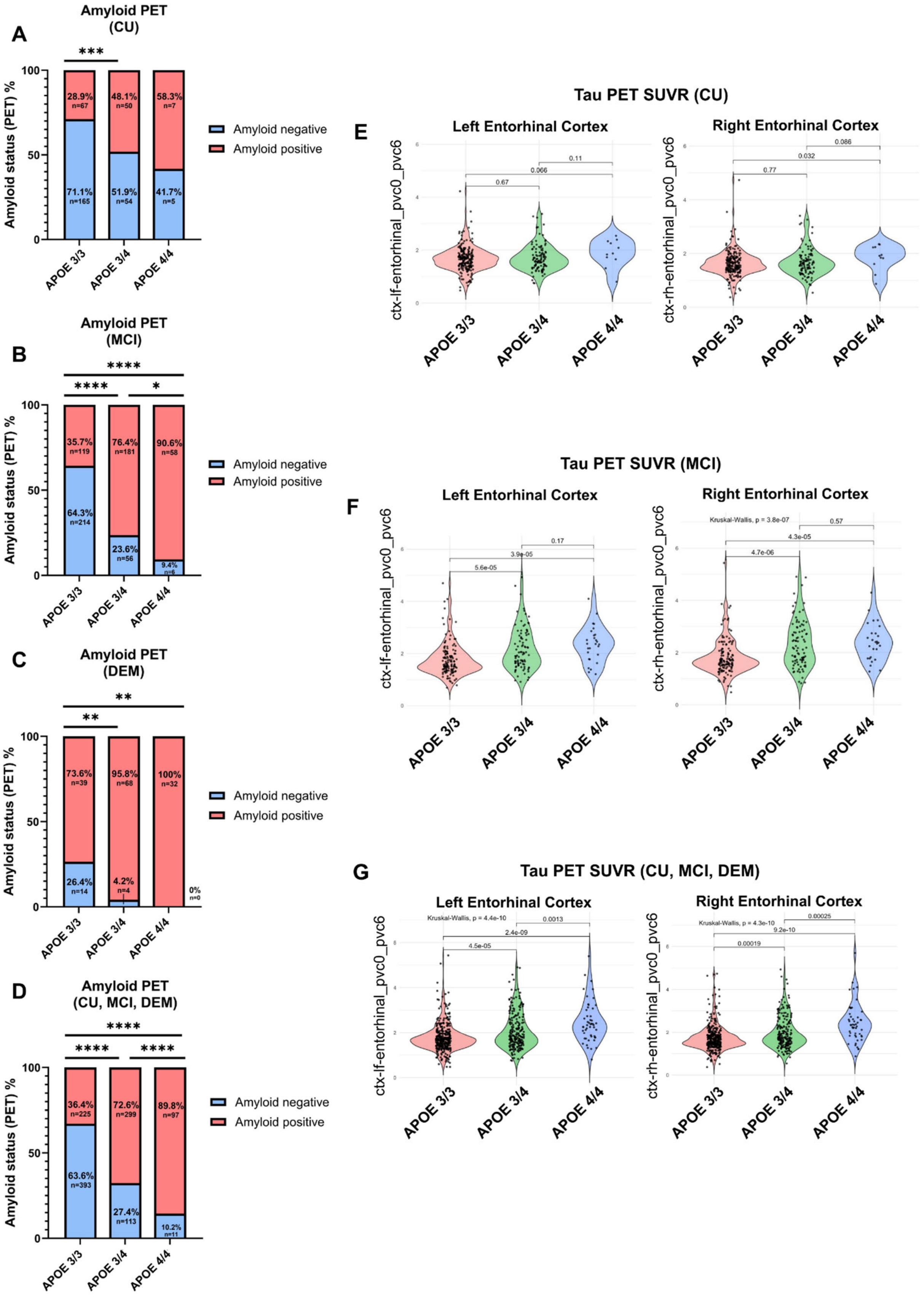

A
Amyloid PET (CU)
***
Amyloid status (PET) %
28.9% n=67
48.1% n=50
58.3% n=7
71.1% n=165
51.9% n=54
41.7% n=5
APOE 3/3
APOE 3/4
APOE 4/4
Amyloid negative
Amyloid positive
B
Amyloid PET (MCI)
****
****
*
35.7% n=119
76.4% n=181
90.6% n=58
64.3% n=214
23.6% n=56
9.4% n=6
C
Amyloid PET (DEM)
**
**
73.6% n=39
95.8% n=68
100% n=32
26.4% n=14
4.2% n=4
0% n=0
D
Amyloid PET (CU, MCI, DEM)
****
****
****
36.4% n=225
72.6% n=299
89.8% n=97
63.6% n=393
27.4% n=113
10.2% n=11
E
Tau PET SUVR (CU)
Left Entorhinal Cortex
Right Entorhinal Cortex
ctx-lf-entorhinal_pvc0_pvc6
ctx-rh-entorhinal_pvc0_pvc6
0.67
0.066
0.11
0.77
0.032
0.086
F
Tau PET SUVR (MCI)
Kruskal-Wallis, p = 3.8e-07
5.6e-05
3.9e-05
0.17
4.7e-06
4.3e-05
0.57
G
Tau PET SUVR (CU, MCI, DEM)
Kruskal-Wallis, p = 4.4e-10
4.5e-05
2.4e-09
0.0013
Kruskal-Wallis, p = 4.3e-10
0.00019
9.2e-10
0.00025

**Figure S1. APOE4 status significantly increases amyloid prevalence and tau burden by ADNI PET data, related to Figure 1.**

**A.** Quantification of proportion of population with positive amyloid PET scan by APOE genotype in CU cohort. $n$ = 232, 104, and 12 for APOE3/3, APOE3/4, and APOE4/4 respectively.

**B.** Quantification of proportion of population with positive amyloid PET scan by APOE genotype in MCI cohort. $n$ = 333, 237, and 64 for APOE3/3, APOE3/4, and APOE4/4 respectively.

**C.** Quantification of proportion of population with positive amyloid PET scan by APOE genotype in DEM cohort. $n$ = 53, 71, and 32 for APOE3/3, APOE3/4, and APOE4/4 respectively.

**D.** Quantification of proportion of population with positive amyloid PET scan by APOE genotype in total population (CU, MCI, and DEM). $n$ = 618, 412, and 108 for APOE3/3, APOE3/4, and APOE4/4 respectively.

**E.** Quantification of normalized Tau PET SUVR in left and right entorhinal cortex by APOE genotype in CU cohort. $n$ = 205, 106, and 11 for APOE3/3, APOE3/4, and APOE4/4 respectively.

**F.** Quantification of normalized Tau PET SUVR in left and right entorhinal cortex by APOE genotype in MCI cohort. $n$ = 134, 90, and 27 for APOE3/3, APOE3/4, and APOE4/4 respectively.

**G.** Quantification of normalized Tau PET SUVR in left and right entorhinal cortex by APOE genotype in total population. $n$ = 17, 15, and 12 for APOE3/3, APOE3/4, and APOE4/4 respectively.

* = $p < 0.05$, ** = $p < 0.01$, *** = $p < 0.001$, **** = $p < 0.0001$. Amyloid PET statistical analysis was done by Chi-square Test with Yates' correction (A, B, C, and D). A positive amyloid scan is defined by a normalized SUVR > 1.11 (see methods). Tau PET statistical analysis was done by Kruskal-Wallis test (E, F, and G) with $p$ values shown.

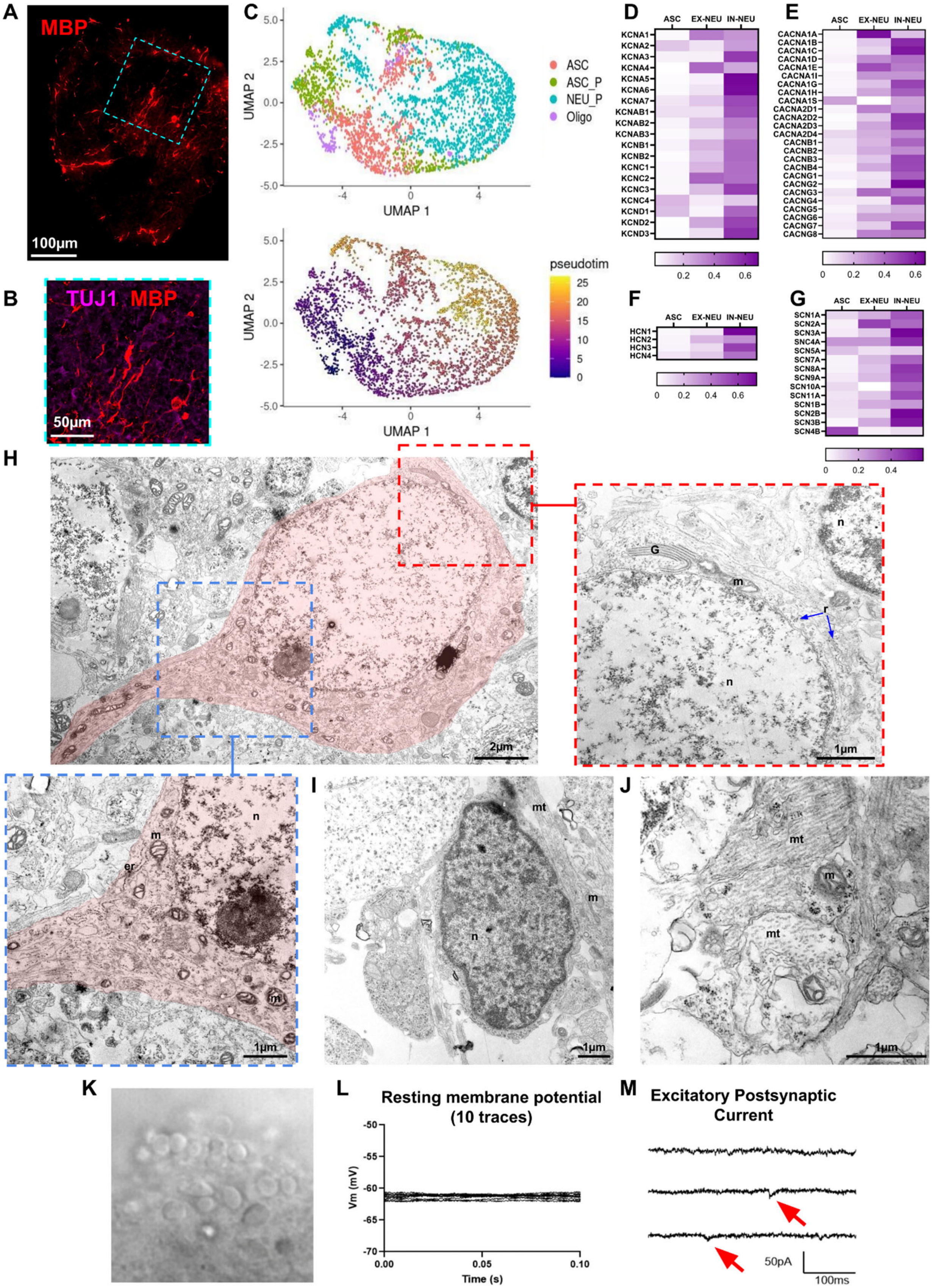
A
MBP
100μm
B
TUJ1 MBP
50μm
C
UMAP 2
UMAP 1
ASC
ASC_P
NEU_P
Oligo
pseudotim
D
ASC EX-NEU IN-NEU
KCNA1
KCNA2
KCNA3
KCNA4
KCNA5
KCNA6
KCNA7
KCNAB1
KCNAB2
KCNAB3
KCNB1
KCNB2
KCNC1
KCNC2
KCNC3
KCNC4
KCND1
KCND2
KCND3
E
CACNA1A
CACNA1B
CACNA1C
CACNA1D
CACNA1E
CACNA1I
CACNA1G
CACNA1H
CACNA1S
CACNA2D1
CACNA2D2
CACNA2D3
CACNA2D4
CACNB1
CACNB2
CACNB3
CACNB4
CACNG1
CACNG2
CACNG3
CACNG4
CACNG5
CACNG6
CACNG7
CACNG8
F
HCN1
HCN2
HCN3
HCN4
G
SCN1A
SCN2A
SCN3A
SNC4A
SCN5A
SCN7A
SCN8A
SCN9A
SCN10A
SCN11A
SCN1B
SCN2B
SCN3B
SCN4B
H
2μm
1μm
G
m
n
r
er
I
mt
J
K
L
Resting membrane potential (10 traces)
Vm (mV)
Time (s)
M
Excitatory Postsynaptic Current
50pA
100ms

**Figure S2. Masteroids exhibit structural and biophysical properties reminiscent of the human brain, related to Figure 2.**

**A.** Representative tilescan image of oligodendrocytes in 14μm thin section of APOE4 NC Aβ Masteroid by MBP (red). Scale bar = 100μm.

**B.** Representative z-stack image of higher magnification of oligodendrocytes in 14μm thin section of whole APOE4 NC Aβ Masteroid by MBP (red). Neurons are also shown (TUJ1, magenta). Scale bar = 50μm.

**C.** scRNA-sequencing pseudotime trajectory analysis of oligodendrocytes in Masteroids. Top UMAP is a reclustering of neuron progenitor cells (NEU_P, cyan), astrocyte progenitor cells (ASC_P, green), astrocytes (ASC, red), and oligodendrocytes (Oligo, purple). Bottom UMAP is the pseudotime analysis of clustered progenitors, astrocytes, and oligodendrocytes.

**D.** scRNA-seq relative gene expression of voltage-gated potassium channel subunits.

**E.** scRNA-seq relative gene expression of voltage-gated calcium channel subunits.

**F.** scRNA-seq relative gene expression of hyperpolarization-activated cyclic nucleotide-gated (HCN) channel subunits.

**G.** scRNA-seq relative gene expression of voltage-gated sodium channel subunits

**H.** Electron microscopic (EM) image of a neuron and neurite (red highlight). Top left image is a combination of 2 overlapping images. The blue-outlined image shows a magnified view of the cytoplasmic area of a neuron and an extending neurite. The red-outlined image shows a magnified view of the perinuclear cytoplasmic area of a neuron adjacent to a glia cell. Blue arrows point to ribosomes. Scale bar = 2μm (top left), 1μm (red and blue outline).

**I.** EM image of a glia cell. Scale bar = 1μm.

**J.** EM image of microtubules within neurites. Scale bar = 1μm

**K.** DIC image of Masteroid used for whole-cell patch-clamp single cell recording.

**L.** Representative traces (10 traces) of resting membrane potential of hiNeurons in Masteroid.

**M.** Representative trace of excitatory postsynaptic current (EPSCs) in Masteroid. Red arrows point to examples of EPSC.

(H-J) m: mitochondria, n: nucleus, er: endoplasmic reticulum, G: Golgi, mt: microtubules, r: ribosomes.

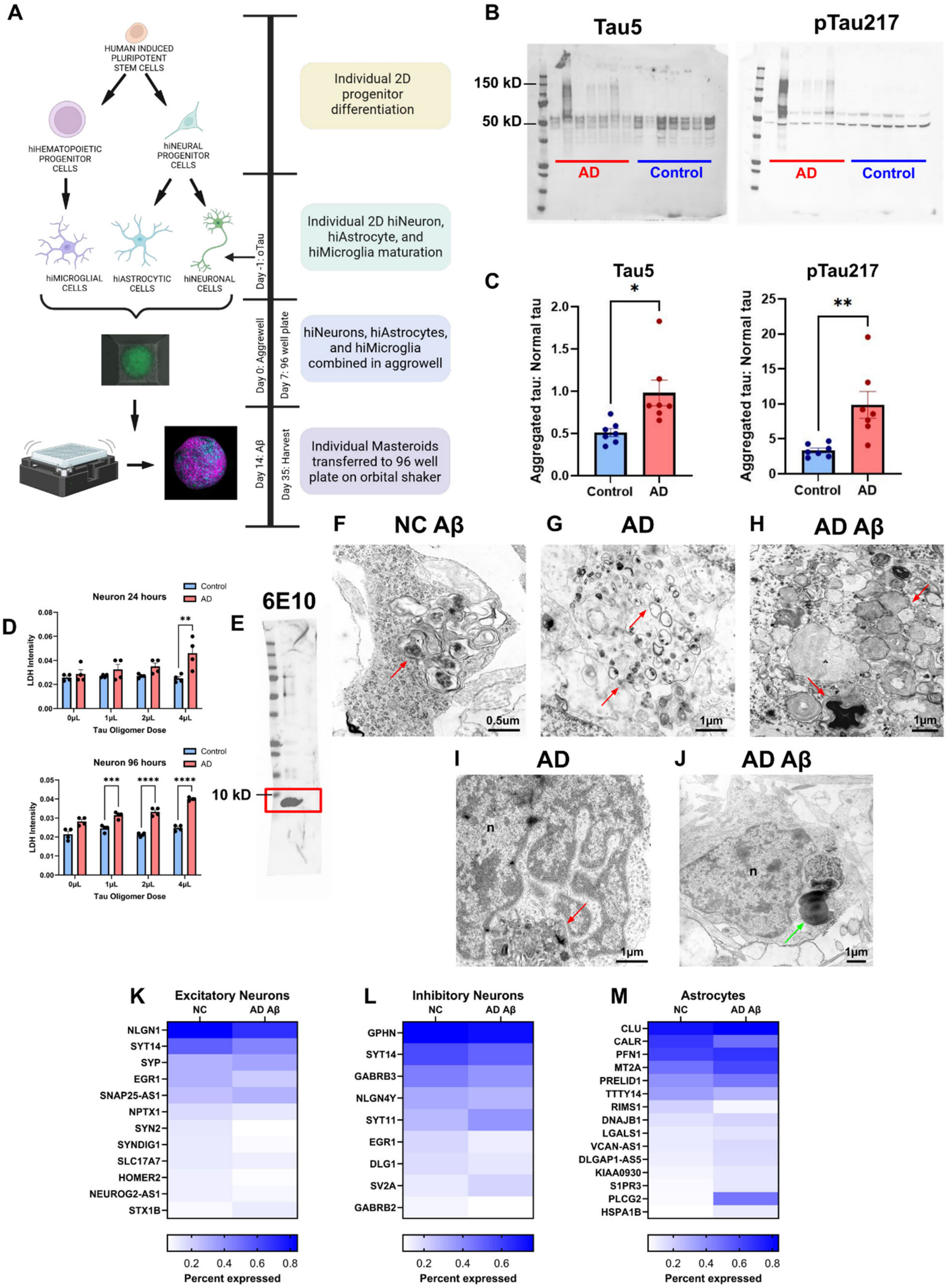
A
HUMAN INDUCED PLURIPOTENT STEM CELLS
hiHEMATOPOIETIC PROGENITOR CELLS
hiNEURAL PROGENITOR CELLS
hiMICROGLIAL CELLS
hiASTROCYTIC CELLS
hiNEURONAL CELLS
Day -1: oTau
Day 0: Aggrewell
Day 7: 96 well plate
Day 14: Aβ
Day 35: Harvest
Individual 2D progenitor differentiation
Individual 2D hiNeuron, hiAstrocyte, and hiMicroglia maturation
hiNeurons, hiAstrocytes, and hiMicroglia combined in aggrowell
Individual Masteroids transferred to 96 well plate on orbital shaker
B
Tau5
pTau217
150 kD
50 kD
AD
Control
C
Aggregated tau: Normal tau
Control
AD
D
Neuron 24 hours
Neuron 96 hours
Control
AD
LDH Intensity
Tau Oligomer Dose
0μL
1μL
2μL
4μL
E
6E10
10 kD
F
NC Aβ
0.5um
G
AD
1μm
H
AD Aβ
1μm
I
AD
n
1μm
J
AD Aβ
n
1μm
K
Excitatory Neurons
NC
AD Aβ
NLGN1
SYT14
SYP
EGR1
SNAP25-AS1
NPTX1
SYN2
SYNDIG1
SLC17A7
HOMER2
NEUROG2-AS1
STX1B
L
Inhibitory Neurons
NC
AD Aβ
GPHN
SYT14
GABRB3
NLGN4Y
SYT11
EGR1
DLG1
SV2A
GABRB2
M
Astrocytes
NC
AD Aβ
CLU
CALR
PFN1
MT2A
PRELID1
TTTY14
RIMS1
DNAJB1
LGALS1
VCAN-AS1
DLGAP1-AS5
KIAA0930
S1PR3
PLCG2
HSPA1B
Percent expressed
0.2
0.4
0.6
0.8

**Figure S3. Modification of Masteroid culture system creates AD model, related to Figure 3.**

**A.** Modified Masteroid culture schematic including the addition of S1p tau oligomers to neurons at day -1 and Aβ oligomers at day 14.

**B.** Western blots of S1p tau fraction with Tau5 antibody (left) and pTau217 antibody (right). Samples underlined in red and denoted with "AD" are from diseased post-mortem human brain tissue, and samples underlined in blue and denoted with "Control" (abbreviated "NC") are from healthy age-matched control post-mortem human brain tissue.

**C.** Quantification of misfolded tau normalized to total tau in the S1p Western blots by Tau5 (total tau) and pTau217 (tau phosphorylated at serine 217).

**D.** Dose response of S1p tau fraction in iPSC-derived neurons at 24 (left) and 96 (right) hours by mean LDH intensity.

**E.** Western blot of 1-16 Aβ oligomers probed by 6E10 antibody.

**F.** Electron microscopic (EM) image of degenerative pathology in NC Aβ Masteroid. Red arrows point to examples of dystrophic neurites. Scale bar = 0.5μm.

**G.** EM image of dystrophic neurites and neurodegenerative pathology in AD oTau Masteroid. Red arrows point to examples of dystrophic neurites. Scale bar = 1μm.

**H.** EM image of degeneration including multilaminar structures at various electron densities in AD Aβ Masteroid. Red arrows point to examples of degenerating myelin and membrane sheath disintegration. Scale bar = 1μm.

**I.** EM image of glial engulfment of dystrophic neurites in AD oTau Masteroid. Red arrow points to engulfed dystrophy in glial cytoplasm. Scale bar = 1μm.

**J.** EM image of lipofuscin in a glial cell within an AD Aβ Masteroid. Green arrow points to lipofuscin in glial cytoplasm. Scale bar = 1μm.

**K.** Heatmap of scRNA-seq expression of synaptic genes in excitatory neurons (EX_NEU) in NC and AD Aβ Masteroids.

**L.** Heatmap of scRNA-seq expression of synaptic genes in inhibitory neurons (IN_NEU) in NC and AD Aβ Masteroids.

**M.** Heatmap of scRNA-seq expression of reactive astrocyte genes in astrocytes (ASC) in NC and AD Aβ Masteroids.

(I, J) n: nucleus. Error bars indicate 95% standard error of the mean. * = $p < 0.05$, ** = $p < 0.01$, *** = $p < 0.001$, **** = $p < 0.0001$. Statistical analysis was done with unpaired t-test (C) and two-way ANOVA with Tukey's multiple comparisons test (D).

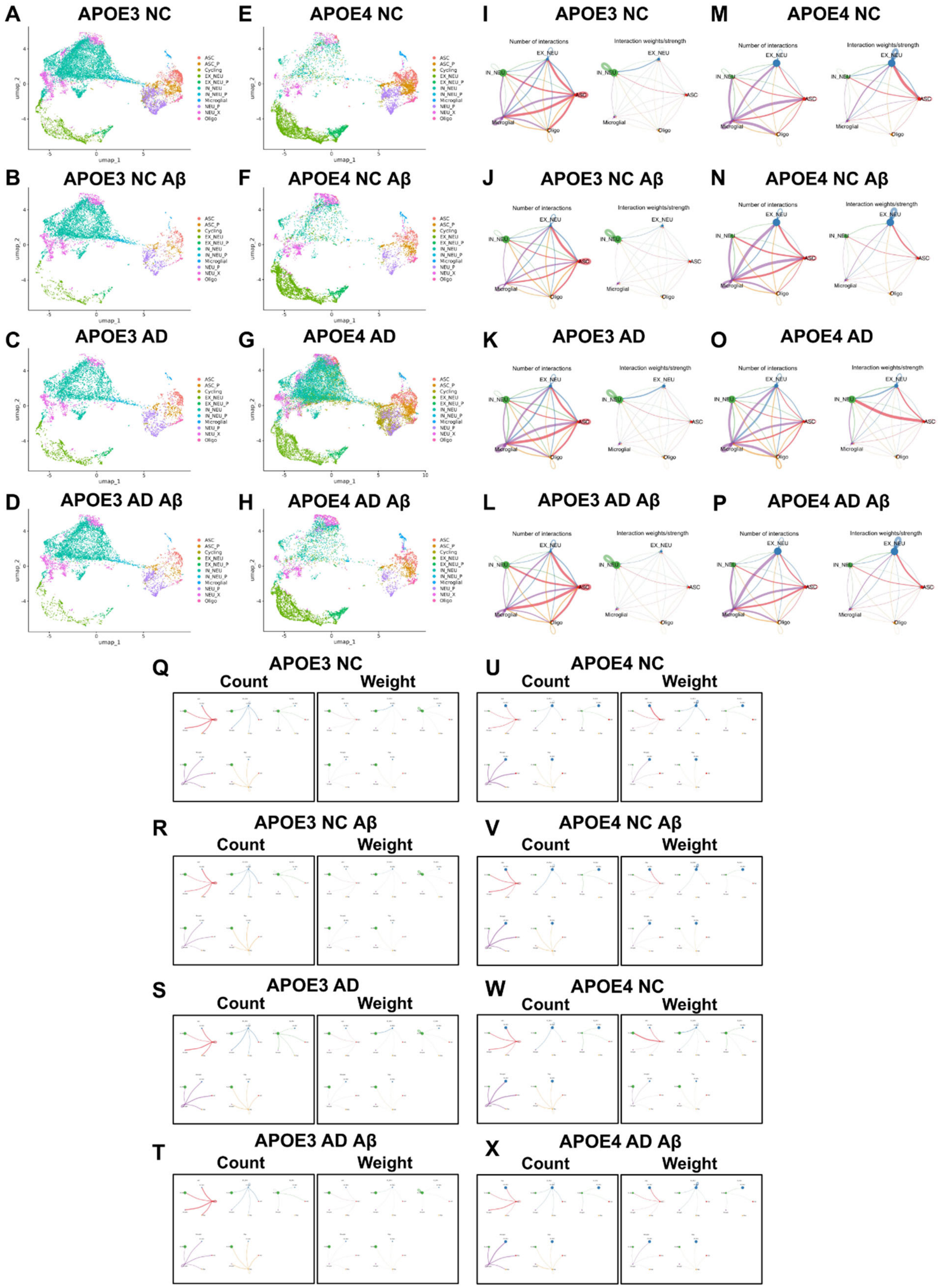
A
APOE3 NC
B
APOE3 NC Aβ
C
APOE3 AD
D
APOE3 AD Aβ
E
APOE4 NC
F
APOE4 NC Aβ
G
APOE4 AD
H
APOE4 AD Aβ
I
APOE3 NC
J
APOE3 NC Aβ
K
APOE3 AD
L
APOE3 AD Aβ
M
APOE4 NC
N
APOE4 NC Aβ
O
APOE4 AD
P
APOE4 AD Aβ
Number of interactions
Interaction weights/strength
Q
APOE3 NC
R
APOE3 NC Aβ
S
APOE3 AD
T
APOE3 AD Aβ
U
APOE4 NC
V
APOE4 NC Aβ
W
APOE4 NC
X
APOE4 AD Aβ
Count
Weight

**Figure S4. scRNA UMAPs and CellChat networks for all Masteroid conditions, related to Figures 2, 3, 4, 5, and 6.**

**A-H.** The following cell types were identified: astrocytes (ASC, red), excitatory neurons (EX_NEU, lime green), inhibitory neurons (IN_NEU, teal), microglia (Microglial, blue), oligodendrocytes (Oligo, pink), and various progenitor and pan-neuronal cells (ASC_P, orange; Cycling, olive; EX_NEU_P, green; IN_NEU_P, cyan; NEU_P, purple; NEU_X, light pink).

**A.** UMAP of APOE3 NC condition.

**B.** UMAP of APOE3 NC Aβ condition.

**C.** UMAP of APOE3 AD oTau condition.

**D.** UMAP of APOE3 AD AB condition.

**E.** UMAP of APOE4 NC condition.

**F.** UMAP of APOE4 NC Aβ condition.

**G.** UMAP of APOE4 AD oTau condition.

**H.** UMAP of APOE4 AD AB condition.

**I-X.** Aggregated networks of number and weight/strength of significant ligand-receptor interactions between the five major cell types. Circle sizes are proportional to the number of cells within a given cluster, and line width is proportional to the indicated number or strength of interactions.

**I.** Aggregated networks of APOE3 NC Masteroids.

**J.** Aggregate networks of APOE3 NC Aβ Masteroids.

**K.** Aggregated networks of APOE3 AD oTau Masteroids.

**L.** Aggregated networks of APOE3 AD Aβ Masteroids.

**M.** Aggregated networks of APOE4 NC Masteroids.

**N.** Aggregate networks of APOE4 NC Aβ Masteroids.

**O.** Aggregated networks of APOE4 AD oTau Masteroids.

**P.** Aggregated networks of APOE4 AD Aβ Masteroids.

**Q.** Cluster specific networks of APOE3 NC Masteroids.

**R.** Cluster specific networks of APOE3 NC Aβ Masteroids.

**S.** Cluster specific networks of APOE3 AD oTau Masteroids.

**T.** Cluster specific networks of APOE3 AD Aβ Masteroids.

**U.** Cluster specific networks of APOE4 NC Masteroids.

**V.** Cluster specific networks of APOE4 NC Aβ Masteroids.

**W.** Cluster specific networks of APOE4 AD oTau Masteroids.

**X.** Cluster specific networks of APOE4 AD Aβ Masteroids.

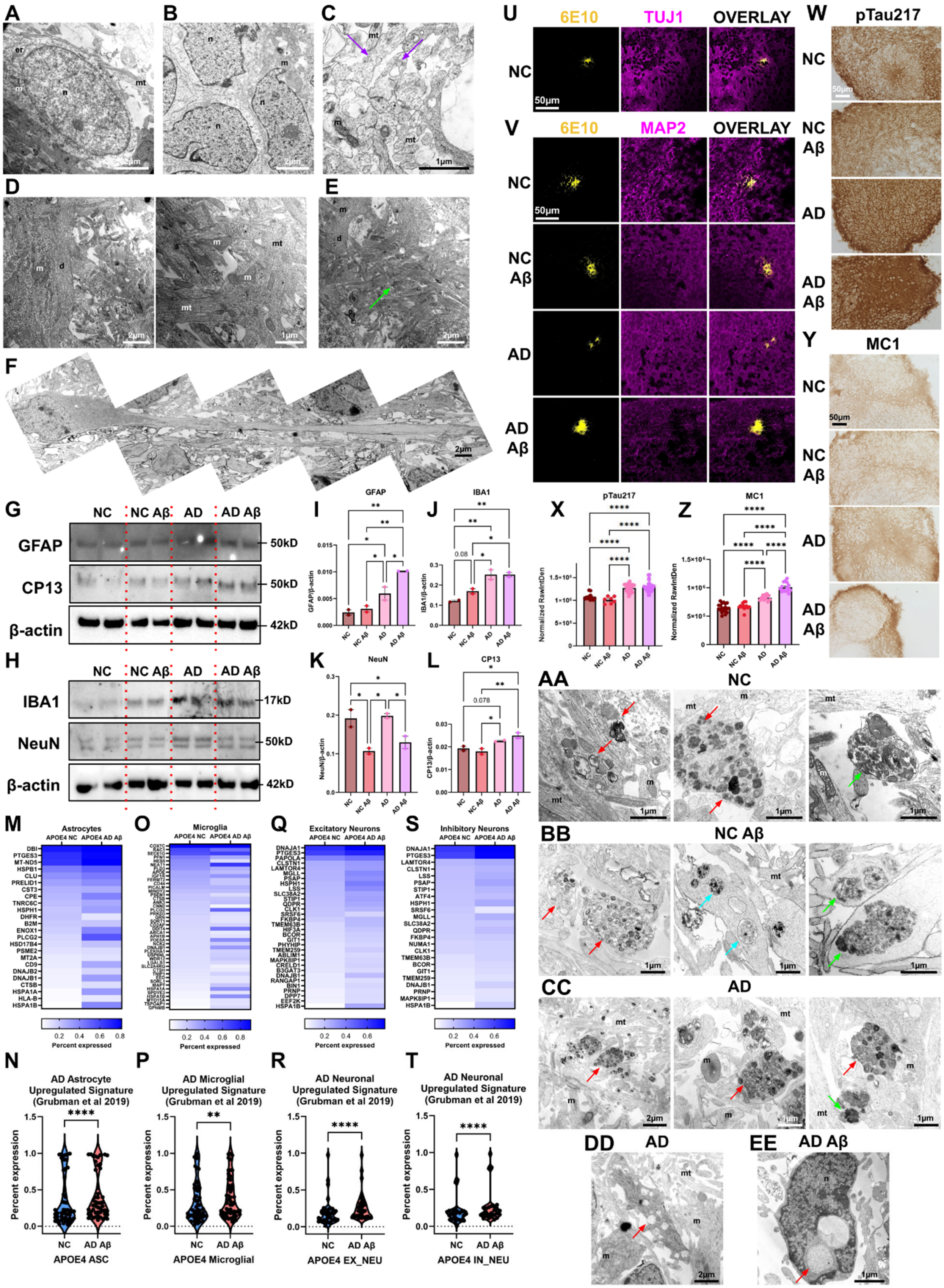
A
B
C
D
E
F
G
NC
NC Aβ
AD
AD Aβ
GFAP
CP13
β-actin
50kD
50kD
42kD
H
IBA1
NeuN
17kD
50kD
42kD
I
GFAP
GFAP/β-actin
J
IBA1
IBA1/β-actin
K
NeuN
NeuN/β-actin
L
CP13
CP13/β-actin
U
6E10
TUJ1
OVERLAY
50μm
V
MAP2
W
pTau217
X
Normalized RawIntDen
Y
MC1
Z
AA
BB
CC
DD
EE
M
Astrocytes
APOE4 NC
APOE4 AD Aβ
O
Microglia
Q
Excitatory Neurons
S
Inhibitory Neurons
Percent expressed
N
AD Astrocyte Upregulated Signature (Grubman et al 2019)
APOE4 ASC
P
AD Microglial Upregulated Signature (Grubman et al 2019)
APOE4 Microglial
R
AD Neuronal Upregulated Signature (Grubman et al 2019)
APOE4 EX_NEU
T
AD Neuronal Upregulated Signature (Grubman et al 2019)
APOE4 IN_NEU
Percent expression

**Figure S5. APOE4 Masteroids exhibit expansive AD pathology, related to Figure 4.**

**A.** Electron microscopic (EM) image of a neuron in APOE4 NC Masteroid. Scale bar = 2μm.

**B.** EM image of glial cells in APOE4 NC Masteroid. Scale bar = 2μm.

**C.** EM image of synaptic vesicles and cross sections of microtubules in APOE4 NC Masteroid. Purple arrows point to examples of vesicles. Scale bar = 1μm.

**D.** EM images of neurites and dendrites in APOE4 NC Masteroid. Scale bar = 2μm (left) and 1 μm (right).

**E**. EM image of cytoskeletal structure and neurites. Green arrow points to cytoskeletal bundles. Scale bar = 2μm.

**F.** EM image of soma and projection in APOE4 NC Aβ Masteroid. Figure is compiled of 5 overlaid images. Scale bar = 2μm.

**G.** Western blot bands of GFAP, CP13, and corresponding β-actin from APOE4 Masteroids. (**Note*: part of the blot from Western blot image in Figure S7H)

**H.** Western blot bands of IBA1, NeuN, and corresponding β-actin from APOE4 Masteroids. (**Note*: part of the blot from the Western blot image in Figures 7E and S7G)

**I**. Quantification of GFAP expression by Western blot in APOE4 Masteroids as shown in *G*, with background subtracted and normalized to β-actin.

**J**. Quantification of IBA1 expression by Western blot in APOE4 Masteroids as shown in *H*, with background subtracted and normalized to β-actin.

**K**. Quantification of NeuN expression by Western blot in APOE4 Masteroids as shown in *H,* with background subtracted and normalized to β-actin.

**L.** Quantification of CP13 expression by Western blot in APOE4 Masteroids as shown in *G,* with background subtracted and normalized to β-actin.

**M.** Heatmap showing scRNA-seq expression of AD-associated and reactive astrocyte genes in astrocytes (ASC) from APOE4 NC and APOE4 AD Aβ Masteroids.

**N.** Comparison of AD-associated astrocyte gene signatures between APOE4 NC and APOE4 AD Aβ astrocytes (ASC) by cross-reference to the human AD snRNA-seq dataset.

**O.** Heatmap showing scRNA-seq expression of AD-associated genes in microglia (Microglial) in APOE4 NC and APOE4 AD Aβ Masteroids.

**P.** Comparison of AD-associated microglial gene signatures between APOE4 NC and APOE4 AD Aβ microglia (Microglial) by scRNA-seq with cross-reference to the human AD snRNA-seq dataset.

**Q.** Heatmap of scRNA-seq expression of AD-upregulated genes identified from human snRNA-seq in excitatory neurons (EX_NEU) from APOE4 NC and APOE4 AD Aβ Masteroids.

**R.** Comparison of AD-associated neuronal signatures in excitatory neurons (EX_NEU) between APOE4 NC and APOE4 AD Aβ Masteroids by scRNA-seq with cross-reference to the human AD snRNA-seq dataset.

**S.** Heatmap of scRNA-seq expression of AD-upregulated genes identified from human snRNA-seq in inhibitory neurons (IN_NEU) from APOE4 NC and APOE4 AD Aβ Masteroids.

**T.** Comparison of AD-associated neuronal signatures in inhibitory neurons (IN_NEU) between APOE4 NC and APOE4 AD Aβ Masteroids by scRNA-seq with cross-reference to the human AD snRNA-seq dataset.

**U.** Representative image of Aβ plaque (6E10, yellow), in a whole APOE4 NC Masteroid. Scale bar = 50μm.

**V.** Representative images of Aβ plaques (6E10, yellow) in 14μm thin sections of APOE4 Masteroids. Scale bar = 50μm.

**W.** Representative DAB images of pTau217 in 14μm thin sections of APOE4 Masteroids. Scale bar = 50μm.

**X.** Quantification of phosphorylated tau in APOE4 Masteroids by pTau217 staining of raw integrated density normalized to Masteroid area.

**Y.** Representative DAB images of MC1 DAB staining in 14μm thin sections of APOE4 Masteroids. Scale bar = 50μm.

**Z.** Quantification of misfolded tau in APOE4 Masteroids by MC1 DAB staining, raw integrated density normalized to Masteroid area.

**AA.** EM images of degenerative pathology including dystrophic neurites, degenerating myelin, and lipofuscin in APOE4 NC Masteroids. Red arrows point to dystrophic neurites and neurodegeneration, green arrow points to lipofuscin. Scale bars = 1μm.

**BB.** EM images of degenerative pathology including dystrophic neurites, degenerating myelin, and lipofuscin in APOE4 NC Aβ Masteroids. Red arrows point to dystrophic neurites, cyan arrows point to encapsulated electron dense granules, green arrows point to lipofuscin. Scale bars = 1μm.

**CC.** EM images of degenerative pathology including dystrophic neurites, degenerating myelin, and lipofuscin in APOE4 AD oTau Masteroids. A range of mitochondrial morphology is observed. Red arrows point to dystrophic neurites, green arrow points to lipofuscin. Scale bars = 2μm (right) and 1μm (center, left).

**DD.** EM image of excessive vacuoles within a singular neurite in AD Masteroid. Red arrow points to vacuoles**.** Scale bar = 2μm.

**EE.** EM image of glial engulfment in APOE4 AD Aβ Masteroid. Red arrows point to engulfed inclusions. Scale bar = 1μm.

(A-E, AA-DD) m: mitochondria, n: nucleus, er: endoplasmic reticulum, mt: microtubules. Error bars indicate 95% standard error of the mean. * = $p < 0.05$, ** = $p < 0.01$, *** = $p < 0.001$, **** = $p < 0.0001$. Statistical analysis was done using one-way ANOVA with Fisher's LSD (I, J, K, and L), Tukey's multiple comparisons test (X and Z), and paired one-sided Wilcoxon signed-rank test (N, P, R, and T).

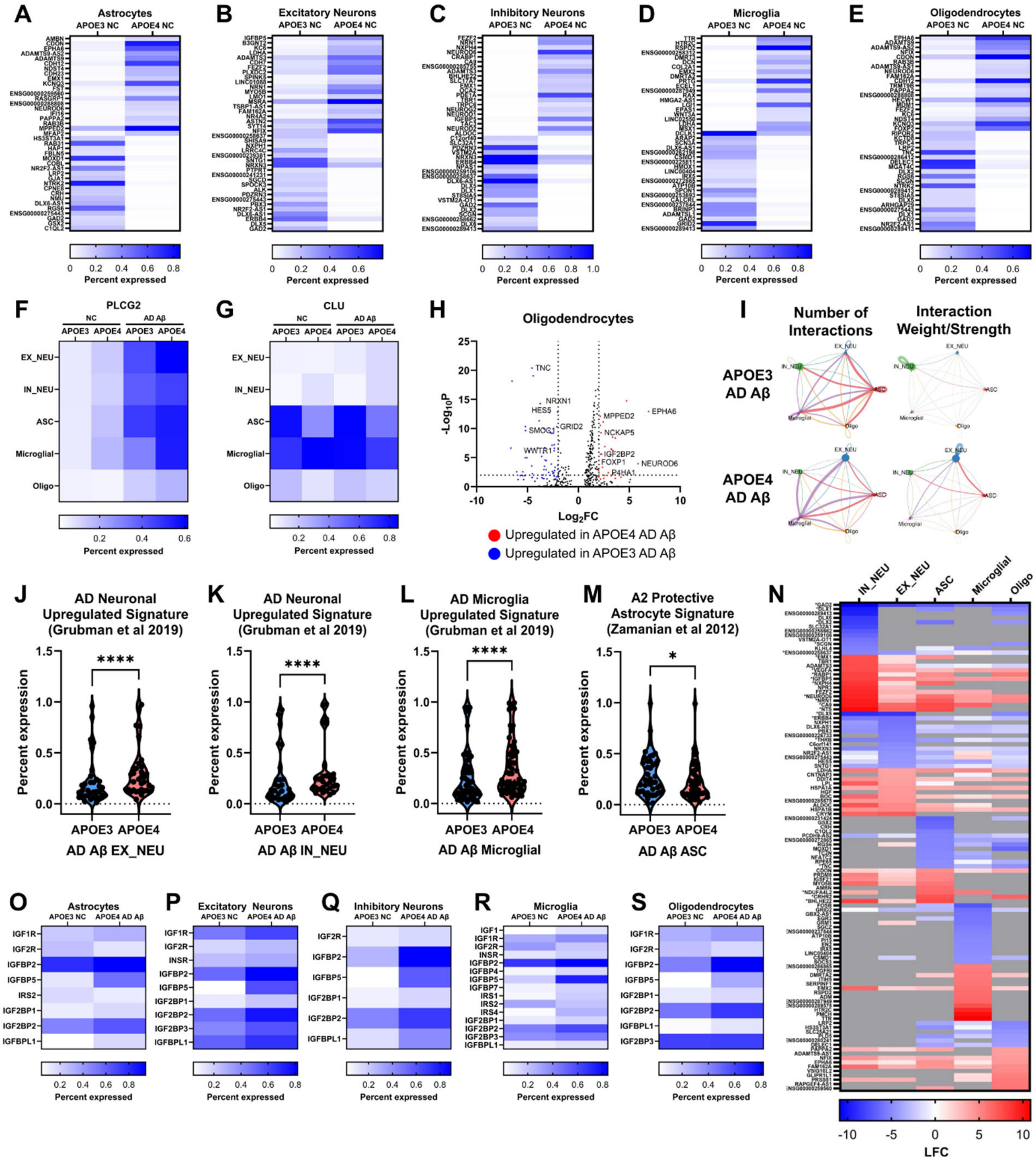

A Astrocytes
B Excitatory Neurons
C Inhibitory Neurons
D Microglia
E Oligodendrocytes
APOE3 NC APOE4 NC
Percent expressed
F PLCG2
G CLU
NC AD Aβ
APOE3 APOE4 APOE3 APOE4
EX_NEU
IN_NEU
ASC
Microglial
Oligo
H Oligodendrocytes
$-Log_{10}P$
$Log_2FC$
TNC
NRXN1
HES5
SMOC1
GRID2
WWTR1
MPPED2
EPHA6
NCKAP5
IGF2BP2
FOXP1
NEUROD6
P4HA1
Upregulated in APOE4 AD Aβ
Upregulated in APOE3 AD Aβ
I Number of Interactions
Interaction Weight/Strength
APOE3 AD Aβ
APOE4 AD Aβ
J AD Neuronal Upregulated Signature (Grubman et al 2019)
K AD Neuronal Upregulated Signature (Grubman et al 2019)
L AD Microglia Upregulated Signature (Grubman et al 2019)
M A2 Protective Astrocyte Signature (Zamanian et al 2012)
Percent expression
****
*
APOE3 APOE4
AD Aβ EX_NEU
AD Aβ IN_NEU
AD Aβ Microglial
AD Aβ ASC
N IN_NEU EX_NEU ASC Microglial Oligo
LFC
O Astrocytes
P Excitatory Neurons
Q Inhibitory Neurons
R Microglia
S Oligodendrocytes
APOE3 NC APOE4 AD Aβ
IGF1
IGF1R
IGF2R
INSR
IGFBP2
IGFBP4
IGFBP5
IGFBP7
IRS1
IRS2
IRS4
IGF2BP1
IGF2BP2
IGF2BP3
IGFBPL1
Percent expressed

**Figure S6. Contribution of APOE4 in driving AD pathogenesis, related to Figures 5 and 6.**

**A-E:** Genes listed in order of most positive to most negative LFC with positive changes representing upregulation in APOE4 Masteroids.

**A.** Percent expression of top 40 differentially expressed genes (DEGs) by LFC (20 positive, 20 negative) between APOE3 NC and APOE4 NC astrocytes (ASC).

**B.** Percent expression of top 40 DEGs by LFC (20 positive, 20 negative) between APOE3 NC and APOE4 NC excitatory neurons (EX_NEU).

**C.** Percent expression of top 40 DEGs by LFC (20 positive, 20 negative) between APOE3 NC and APOE4 NC inhibitory neurons (IN_NEU).

**D.** Percent expression of top 40 DEGs by LFC (20 positive, 20 negative) between APOE3 NC and APOE4 NC microglia (Microglial).

**E.** Percent expression of top 40 DEGs by LFC (20 positive, 20 negative) between APOE3 NC and APOE4 NC oligodendrocytes (Oligo).

**F.** Percent expression of PLCG2 by cell type across genotypes in NC and AD Aβ conditions.

**G.** Percent expression of CLU by cell type across genotypes in NC and AD Aβ conditions.

**H.** Volcano plot of DEGs between APOE3 AD Aβ and APOE4 AD Aβ in oligodendrocytes (Oligo). Red dots indicate genes upregulated in APOE4 AD Aβ, and blue dots indicate genes downregulated in APOE4 AD Aβ.

**I.** Number and weight of significant ligand-receptor interactions between five major cell types between APOE3 AD Aβ vs APOE4 A3 AD Aβ Masteroids. Circle sizes are proportional to the number of cells within a given cluster, and line width is proportional to the indicated number or strength of interactions. Cell types included are excitatory neurons (EX_NEU, blue), inhibitory neurons (IN_NEU, green), astrocytes (ASC, red), microglia (Microglial, purple), and oligodendrocytes (Oligo, orange).

**J.** scRNA-seq gene expression comparison of AD-associated neuronal signatures in excitatory neurons (EX_NEU) in ApoE3 AD Aβ vs ApoE4 AD Aβ Masteroids with cross-reference to the human AD snRNA-seq dataset.

**K.** scRNA-seq gene expression comparison of AD-associated neuronal signatures in inhibitory neurons (IN_NEU) in ApoE3 AD Aβ vs ApoE4 AD Aβ Masteroids with cross-reference to the human AD snRNA-seq dataset.

**L.** scRNA-seq gene expression comparison of AD-associated signatures in microglia (Microglial) in ApoE3 AD Aβ vs ApoE4 AD Aβ Masteroids with cross-reference to the human AD snRNA-seq dataset.

**M.** scRNA-seq gene expression comparison of astrocytes (ASC) in ApoE3 AD Aβ vs ApoE4 AD Aβ Masteroids of A2 protective astrocytic signature.

**N.** Top 30 DEGs by LFC (15 positive, 15 negative) per cell type between APOE3 NC and APOE4 AD Aβ Masteroids. Positive LFC (red) indicates upregulation in APOE4 AD Aβ Masteroids. Genes appearing in multiple cell types among the top LFC are shown only once, grouped with the cell type showing the greatest LFC, and marked with an *. Gray boxes indicate that the comparison was not included in the DEG analysis for that combination.

**O.** Heatmap of scRNA-seq expression of IGF signaling genes in astrocytes (ASC) in APOE3 NC and APOE4 AD Aβ Masteroids.

**P.** Heatmap of scRNA-seq expression of IGF signaling genes in excitatory neurons (EX_NEU) in APOE3 NC and APOE4 AD Aβ Masteroids.

**Q.** Heatmap of scRNA-seq expression of IGF signaling genes in inhibitory neurons (IN_NEU) in APOE3 NC and APOE4 AD Aβ Masteroids.

**R.** Heatmap of scRNA-seq expression of IGF signaling genes in microglia (Microglial) in APOE3 NC and APOE4 AD Aβ Masteroids.

**S.** Heatmap of scRNA-seq expression of IGF signaling genes in oligodendrocytes (Oligo) in APOE3 NC and APOE4 AD Aβ Masteroids.

* = $p < 0.05$, **** = $p < 0.0001$. Statistical analysis was done using one-tailed Wilcoxon matched-pairs signed rank test (J, K, L, and M).

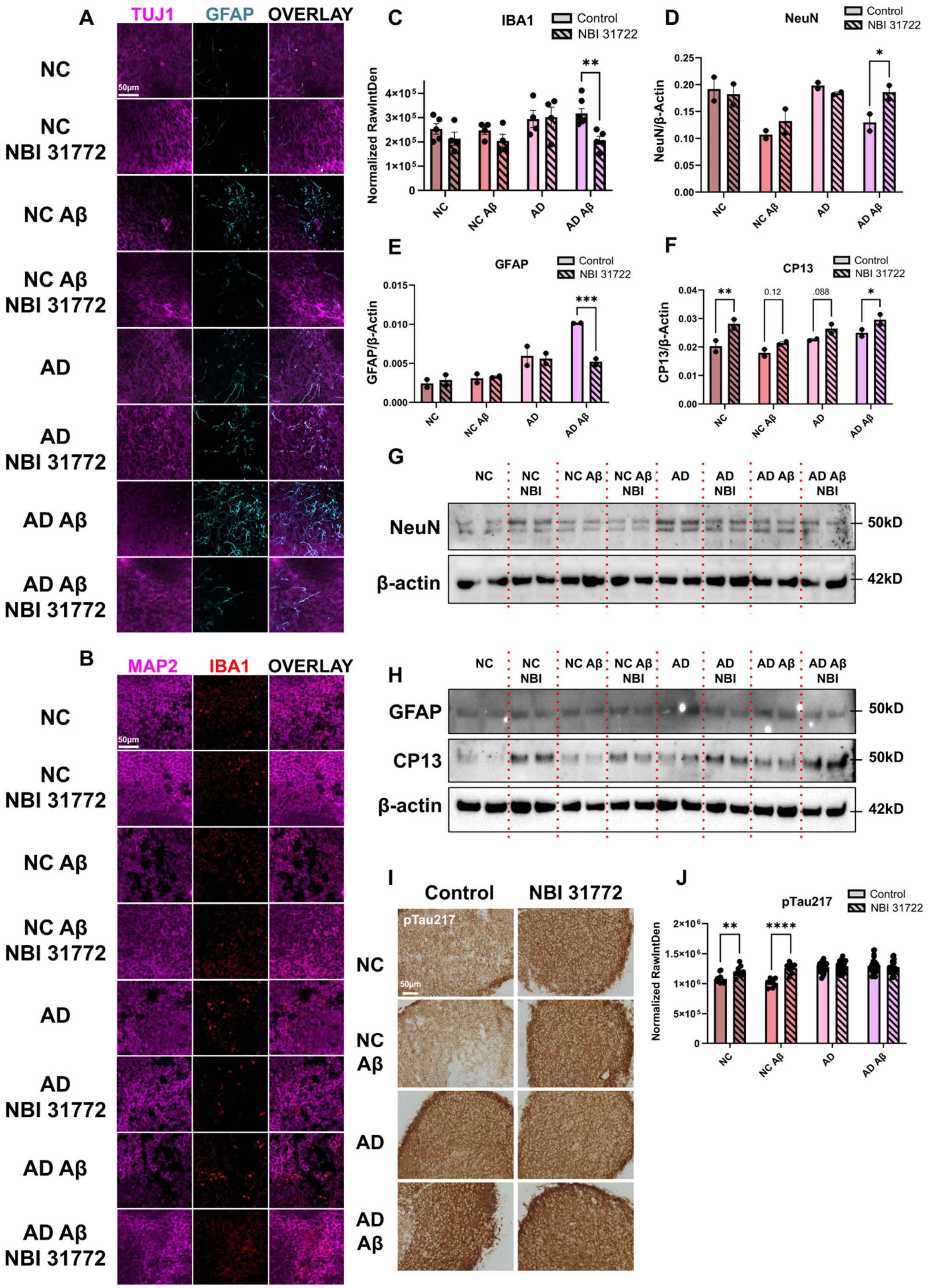

A
TUJ1
GFAP
OVERLAY
NC
NC NBI 31772
NC Aβ
NC Aβ NBI 31772
AD
AD NBI 31772
AD Aβ
AD Aβ NBI 31772
50μm
B
MAP2
IBA1
OVERLAY
C
IBA1
Control
NBI 31722
Normalized RawIntDen
D
NeuN
NeuN/β-Actin
E
GFAP
GFAP/β-Actin
F
CP13
CP13/β-Actin
0.12
.088
G
NeuN
β-actin
50kD
42kD
NC NBI
NC Aβ NBI
AD NBI
AD Aβ NBI
H
GFAP
CP13
I
Control
NBI 31772
pTau217
J

**Figure S7. IGFBP inhibitor NBI 31772 ameliorates AD-associated neuroinflammation, related to Figure 7.**

**A.** High magnification representative immunofluorescence images of APOE4 Masteroids with and without NBI 31772 treatment. Neurons (TUJ1, magenta) and astrocytes (GFAP, cyan) are shown. Scale bar = 50μm.

**B.** High magnification representative immunofluorescence images of 14μm thin sections of APOE4 Masteroid with and without NBI 31772 treatment. Neurons (MAP2, magenta) and microglia (IBA1, red) are shown. Scale bar = 50μm.

**C.** Quantification of microglial activation in NBI 31772 treated APOE4 Masteroids by IBA1 labeling, raw integrated density normalized by Masteroid area.

**D.** Quantification of NeuN expression by Western blot in APOE4 Masteroids with and without NBI 31772 as shown in *G* with background subtracted and normalized to β-actin.

**E.** Quantification of GFAP expression by Western blot in APOE4 Masteroids with and without NBI 31772 as shown in *H* with background subtracted and normalized to β-actin.

**F.** Quantification of CP13 expression by Western blot in APOE4 Masteroids with and without NBI 31772 as shown in *H* with background subtracted and normalized to β-actin.

**G.** Western blot bands of IBA1, NeuN, and corresponding β-actin.

**H.** Western blot bands of GFAP, CP13, and corresponding β-actin.

**I.** Representative images of pTau217 in 14μm thin sections of APOE4 Masteroid with and without NBI 31772 treatment. Scale bar = 50μm.

**J.** Quantification of phosphorylated tau in APOE4 Masteroids with and without NBI 31772 treatment by pTau217 raw integrated density normalized to Masteroid area.

Error bars indicate 95% standard error of the mean. * = $p < 0.05$, ** = $p < 0.01$, *** = $p < 0.001$, **** = $p < 0.0001$. Statistical analysis was done using two-way ANOVA with Fisher's LSD (C, D, E, F, and J).
